# Gene regulatory network topology governs resistance and treatment escape in glioma stem-like cells

**DOI:** 10.1101/2024.02.02.578510

**Authors:** James H. Park, Parvinder Hothi, Adrian Lopez Garcia de Lomana, Min Pan, Rachel Calder, Serdar Turkarslan, Wei-Ju Wu, Hwahyung Lee, Anoop P. Patel, Charles Cobbs, Sui Huang, Nitin S. Baliga

**Affiliations:** Institute for Systems Biology, Seattle, WA; Ivy Center for Advanced Brain Tumor Treatment, Swedish Neuroscience Institute, Seattle, WA; Center for Systems Biology, University of Iceland, Reykjavik, Iceland; Department of Neurosurgery, Preston Robert Tisch Brain Tumor Center, Duke University, Durham, NC; Center for Advanced Genomic Technologies, Duke University, Durham, NC; Departments of Microbiology, Biology, and Molecular Engineering Sciences, University of Washington, Seattle, WA

## Abstract

Poor prognosis and drug resistance in glioblastoma (GBM) manifests from heterogeneity and treatment-induced shifts in phenotypic states of tumor cells, including dedifferentiation to glioma stem-like cells (GSCs). This rare tumorigenic cell subpopulation is inherently resistant to temozolomide, undergoes proneural-to-mesenchymal transition (PMT) to evade therapy, and thereby drives recurrence. Through inference of transcriptional regulatory networks (TRNs) of patient-derived GSCs (PD-GSCs) at single-cell resolution, we demonstrate how topology of transcription factor interactions drives distinct trajectories of cell state transitions of susceptible and resistant PD-GSCs in response to cytotoxic drug treatment. By experimentally testing TRN simulation-based predictions, we show that drug treatment drives surviving cells of a PD-GSC along a trajectory of intermediate states, akin to a bottleneck in gene expression space, exposing vulnerability to potentiated killing by sequential addition of siRNA or a second drug targeting transcriptional programs governing non-genetic plasticity of a PD-GSC. Thus, our findings demonstrate an approach to uncover and use TRN topology of a PD-GSC to rationally predict combinatorial and sequential treatments that block treatment escape and acquired resistance in GBM.

## INTRODUCTION

Glioblastoma (GBM) is the most lethal and aggressive primary brain tumor in adults. With current standard of care (SOC), which involves maximal surgical resection, fractionated radiotherapy (XRT), and chemotherapy with the DNA-alkylating agent, temozolomide (TMZ) (*1*), patient prognosis remains dismal with a median survival time of 14-15 months and a 90% risk of recurrence. There is growing evidence that the poor therapy responsiveness and dismal prognosis in GBM patients emerges from the interplay of tumor cell heterogeneity and treatment-induced shifts of cellular phenotypic states. Three molecular subtypes of GBM have been identified– proneural (PN), classical (CL), and mesenchymal (MES), each exhibiting distinct responses to SOC and clinical prognosis (*2*, *3*). Single-cell resolution transcriptome analyses further demonstrated that even an individual GBM tumor is heterogeneous, not only morphologically but also with respect to its composition of cellular states (*4*), which can include a mixture of PN/CL/MES subtype cells and a small subpopulation of glioma stem-like cells (GSCs) that have the capability to self-renew, generate different tumor cell progenies, and initiate new tumors. Further, there is evidence that extrinsic signals and stressors, including those generated by treatment, can also drive heterogeneous tumor cells to dedifferentiate into immature GSCs that are inherently resistant to TMZ (*5*, *6*).

While PN GSCs have higher proliferation rates and promote tumor angiogenesis, MES GSCs have potent invasive capabilities (*7*) and are more resistant to radiation (*8*) and drug treatment (*9*). Thus, most recurrent tumors derived from non-MES primary tumor are comprised of the MES subtype (*10*, *11*). Two hypotheses have been proposed for the shift in recurrent tumor subtype and corresponding development of treatment resistance (*12*, *13*): 1) MES subtype GSCs are selected for and eventually drive the growth of the recurrent tumor (*14*), or 2) radiation and chemotherapy causes GSCs to undergo a PN to MES transition (PMT) to evade and survive treatment (*7*, *15*). The latter hypothesis is in line with the emerging notion that non-genetic cell plasticity, in addition to selection of fixed, genetically determined phenotypes of mutant cells accounts for tumor progression and recurrence. For instance, radiation– or chemotherapy-induced epithelial to MES transition (EMT) in solid tumors has been widely implicated in the rapid development of therapy resistance (*16–25*). Thus, GSCs undergoing PMT may be causally responsible for recurrence of most drug resistant GBM tumors in the form of the MES subtype (*26*). For example, expression of MES marker (CD44) and NF-kB pathways associated with PMT were elevated following radiation treatment of PN GSCs pretreated with TNF-alpha. In genetically engineered mouse models with cells that can fluorescently report molecular subtype, GSCs transitioned to the MES subtype as early as 6 hours following radiation treatment, demonstrating intrinsic ability of GSCs to deal with treatment-induced stress (*15*). Finally, GSCs isolated from the invasive tumor edge transitioned from a PN subtype to a MES phenotype in a C/EBP-β dependent manner following treatment (*27*). In view of the accumulating evidence for the role of non-genetic plasticity of GSCs in the development of recurrent and refractory tumors, multiple clinical trials are underway to evaluate novel drugs or drug combinations that are both cytotoxic against GSCs and also meet the criteria for treating brain tumors (e.g., penetrance of blood brain barrier) to treat recurrent therapy-refractory GBM (*28*). These clinical studies, including our own, have discovered that many FDA-approved drugs are effective in killing GSCs, but can also induce surviving cells to undergo PMT.

Here, we sought to understand if knowledge of mechanisms of plasticity of GSCs, and the trajectories through which they undergo drug-induced PMT, would enable rational strategies to improve treatment responsiveness by disrupting primary resistance mechanisms, while blocking therapy escape to prevent acquired resistance and tumor recurrence. We have performed these studies with pitavastatin, an HMG-CoA reductase inhibitor, which is widely used to manage cholesterol levels. Pitavastatin is a prime example of an FDA-approved drug that can be repurposed to minimize GBM recurrence because of its anti-proliferative and radiotherapy sensitization effects on glioma cells (*29*) as well as its cytotoxic effects against GSCs (*30*). Specifically, we have investigated mechanisms of primary and acquired resistance in six patient-derived GSCs (PD-GSCs) – three responders (SN520, SN533, and SN575) and three non-responders (SN503, SN517 and SN521) to pitavastatin. Through the inference of mechanistic transcriptional regulatory networks at single cell resolution, we demonstrate that the architecture and dynamics of a core transcription factor (TF) network governed the phenotypic plasticity of PD-GSCs. By performing *in silico* simulations and chemical and genetic (siRNA) perturbations, we show compelling evidence that it wasn’t the composition of initial cell states, but the topology of the core TF-TF network that governed phenotypic plasticity of GSCs. Finally, our findings demonstrate that mechanistic knowledge of the gene regulatory network topology can be leveraged to rationally tailor combinatorial and sequential treatment regimen to disrupt primary or acquired resistance in a given PD-GSC.

## RESULTS

### Pitavastatin treatment induces distinct responses in SN520 and SN503 PD-GSCs

Through high throughput dose titration assays we discovered that pitavastatin had a wide range of effectiveness against 45 PD-GSCs. Based on their varying sensitivities, we classified the PD-GSCs into two categories, one in which PD-GSCs were considered a “responder” (IC50 < 5.0μM) and the other in which they were considered a “non-responder” (IC50 ≥ 5.0μM, Figure 1A). To understand the dynamics underlying each drug-response phenotype, we examined pitavastatin sensitivity of two PD-GSC cultures, SN520 and SN503, both of which were isocitrate dehydrogenase 1 (IDH1) wild-type and O6-methylgaunine-DNA methyltransferase (MGMT) unmethylated. The dose titration results revealed distinct susceptibility profiles to pitavastatin treatment. With an IC_50_ of 13.0μM, SN503 was considered a “non-responder”, whereas as SN520 with an IC50 of 0.43μM was labeled a “responder” (Figure. 1A). Next, we investigated the longitudinal response of each PD-GSC culture over a 4-day treatment with DMSO (vehicle control) or pitavastatin at 6μM, a dose at which significant decreases in cell viability were observed over the same treatment period (Supplementary Figure S1). To minimize batch effects, replicate cultures were treated with drug or vehicle over a staggered schedule such that all samples for days 0 (D0), 2 (D2), 3 (D3), and 4 (D4) were collected and processed simultaneously for subsequent flow cytometry, bulk RNA-seq, and scRNA-seq analysis (Figure. 1B). SN520 viability decreased dramatically during treatment between D3 and D4, falling below 90% by day 5 (Figure 1A). By contrast, over the first three days of pitavastatin treatment, SN503 viability decreased rapidly at a rate that was similar to the kill rate of SN520, but leveled off to ∼60% for the remaining duration of the 4-day treatment.

**Figure 1.**
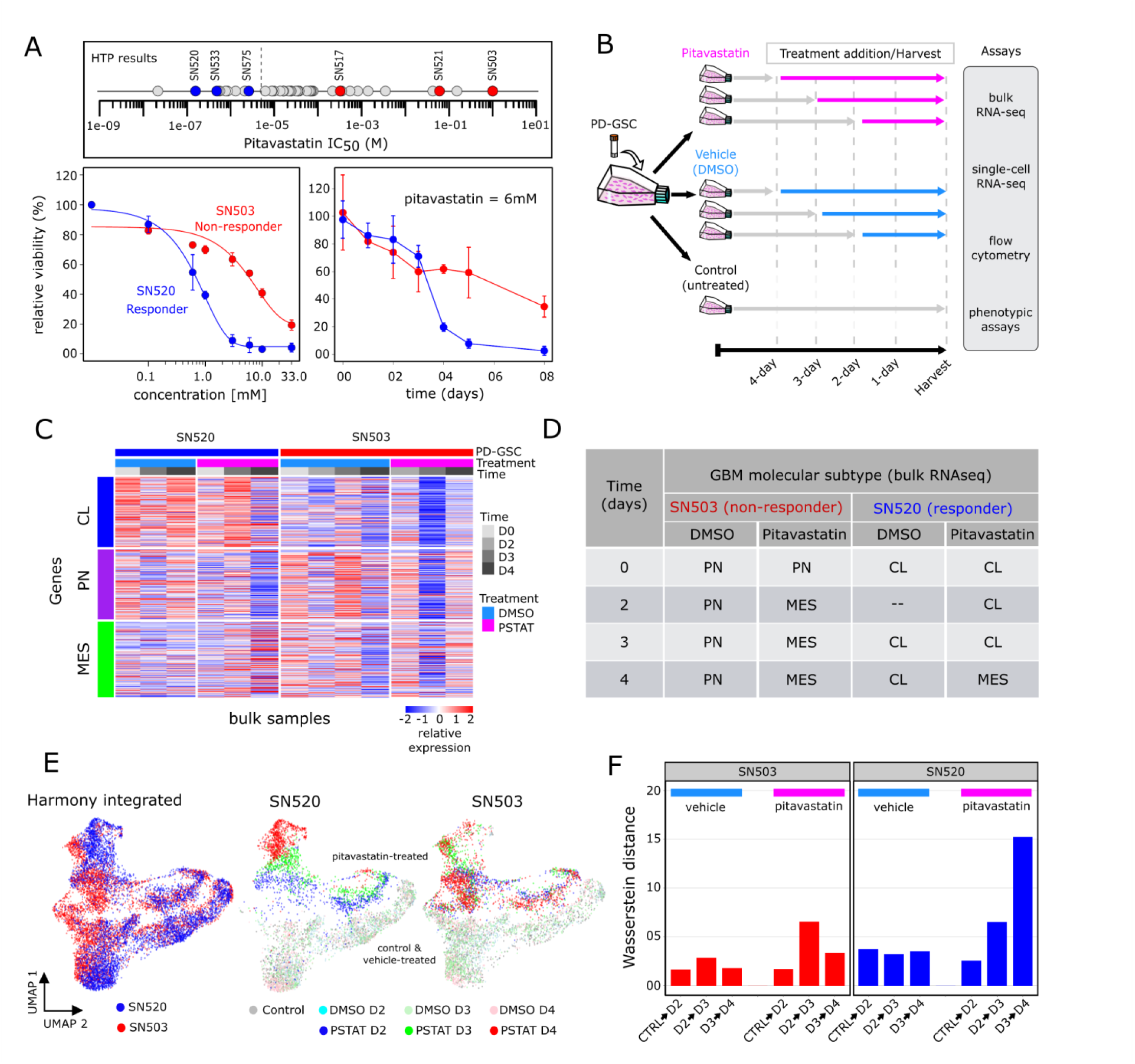
Pitavastatin causes shift in molecular subtype expressed by PD-GSCs. (**A**) Pitavastatin IC_50_ values for each of 45 PD-GSCs as determined using dose titration assays (below). Labeled PD-GSCs represent a subset deemed as a responders (blue) and non-responders (red) to pitavastatin. Below are drug-dose response and time-course response curves for SN520 (pitavastatin-responsive) and SN503 (pitavastatin-non-responsive) PD-GSC populations. **(B)** Experimental workflow for longitudinal monitoring of PD-GSC response to pitavastatin treatment. Colored horizontal arrows indicate duration of pitavastatin (magenta), vehicle-control (DMSO, light blue), or untreated control (dark grey). **(C)** Heatmap of bulk-level expression for molecular subtype gene sets (classical – CL, proneural – PN, mesenchymal – MES) for samples collected. **(D)** Table summarizing dominant molecular subtype expressed in each sample. D2 bulk sample for SN520 was absent due to sample limitations. **(E)** UMAP plots of Harmony-integrated scRNA-seq data sets and corresponding individual plots for each PD-GSC phenotype. **(F)** Wasserstein distance of transport distances between each consecutive time point for each PD-GSC under each treatment condition (vehicle– or pitavastatin-treatment).

Flow cytometry analysis with annexin V labeling demonstrated that pitavastatin had killed SN520 cells by inducing apoptosis (Supplementary Figure S2). This discrepancy was interesting because unlike SN520, cytometry analysis of the SN503 did not reveal any dramatic increase in annexin V signal, suggesting that in this PD-GSC culture a mechanism other than apoptosis was responsible for cell death in a small fraction of the population (Supplementary Figure S2). These findings indicated that the cytotoxic consequences of pitavastatin may vary depending on the composition and characteristics of subpopulations of cells within each PD-GSC culture. Further, the difference in the rate of cell death in both PD-GSC cultures during treatment suggested either the presence of distinct sub-populations of cells with varying susceptibility to pitavastatin, or the induction of adaptive responses and cell state transitions across sub-populations within each PD-GSC culture. In support of this hypothesis, subsequent bulk RNA-seq profiling and gene set variance analysis (GSVA, (*31*)) revealed that while the dominant subtype composition of the two cell cultures was stable during vehicle treatment (DMSO), in response to pitavastatin treatment both PD-GSCs underwent a transition to a MES subtype. While SN503 underwent a rapid shift from PN to MES subtype within two days of treatment, SN520 cells maintained a dominant CL signature for the first three days and then shifted to a MES subtype on the fourth day of treatment (Figure 1C, D). These findings established that despite their similarity in terms of IDH1 mutation and MGMT methylation status, the two PD-GSC cultures exhibited vastly different pitavastatin responses that likely manifested the presence of distinct sub-populations capable of cell state transitions that enabled the surviving cells to escape drug-induced cytotoxicity.

### Single-cell analysis suggests drug-induced PMT is likely mechanism of acquired pitavastatin resistance in SN520

To further dissect the likely role of sub-population heterogeneity in enabling treatment escape of SN520 and SN503 (Figure 1B), we performed scRNA-seq profiling of each PD-GSC culture (Chromium, 10X Genomics, Inc.). Following QC of the raw scRNA-seq data (METHODS), a total of 5,402 cells from SN520 and 5,722 cells from SN503 were profiled across all time points (D0, D2, D3, and D4) and treatment conditions (pitavastatin or vehicle control). Batch-integration with Harmony (*32*), dimensionality reduction, and visualization with uniform manifold approximation and projection (UMAP, (*33*)) of the integrated scRNA-seq data revealed distinct pitavastatin-specific transcriptional responses across the two PD-GSCs (Figure 1E). In SN520, we observed time-dependent clustering of cells, indicating a coordinated transcriptional response to pitavastatin. By contrast, there was considerable overlap between pitavastatin-treated SN503 cells from all time points (Figure 1E). We quantified net temporal shifts in transcriptomic states of the cells, or lack thereof, using Wasserstein distance, which quantifies dissimilarity between two high-dimensional distributions (*34*). Drug treatment caused the SN520 cells to become progressively dissimilar from the preceding state over time, unlike vehicle-treated cells. By contrast, there was a slight increase in Wasserstein distance in drug-treated SN503 cells between D2 and D3, but not between D3 and D4 samples (Figure 1F). Given the distinct response patterns of the two PD-GSCs, subsequent scRNA-seq analysis was performed on a patient-specific basis, (Figure 2A, B). UMAP plots organized cells within each PD-GSC into two main groups, defined by treatment with either pitavastatin or vehicle control. Pitavastatin-treated SN520 cells organized along treatment time whereas pitavastatin-treated SN503 cells from different time points overlapped with one another in the gene expression space as captured by the UMAP embeddings.

**Figure 2.**
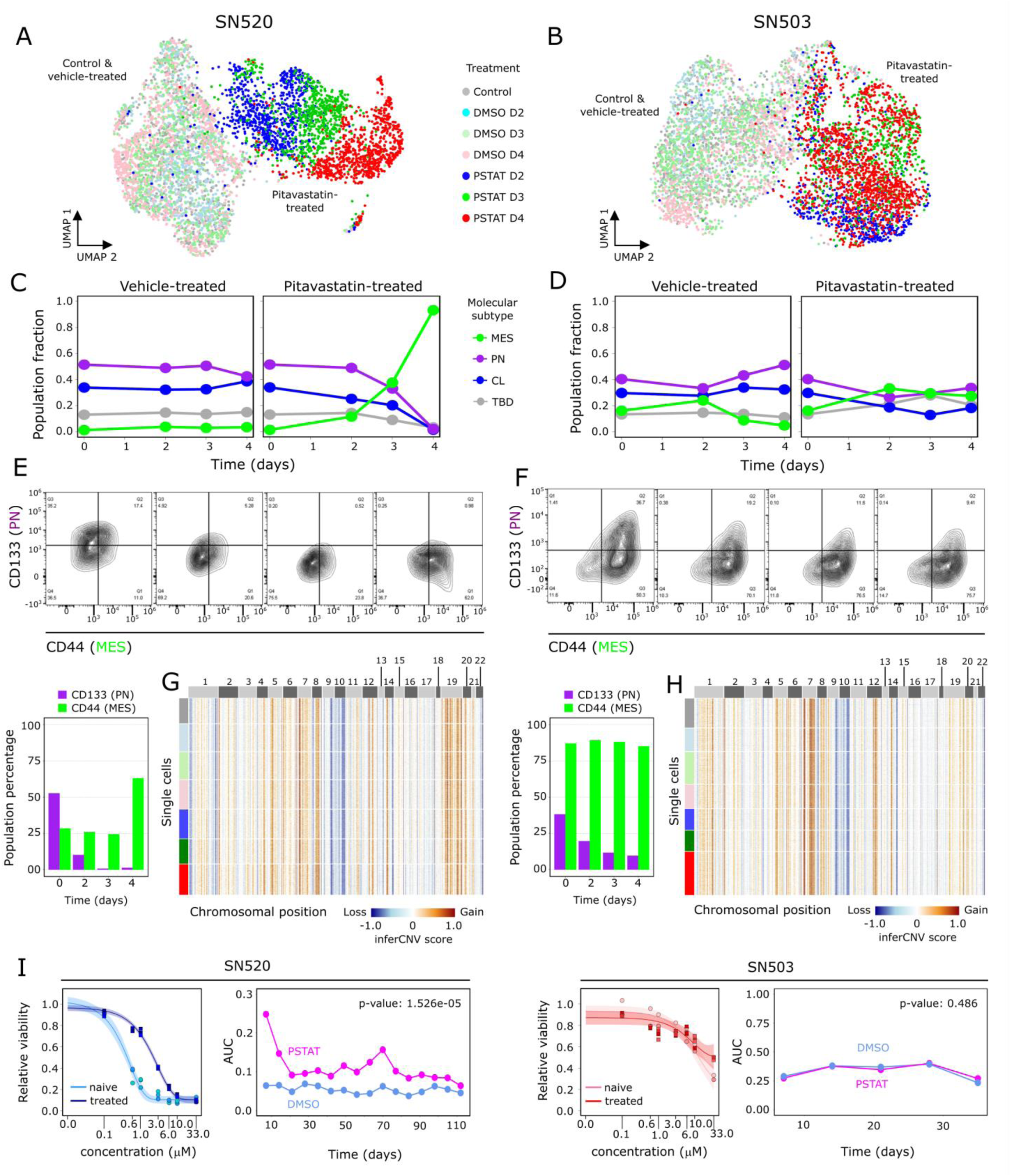
Single-cell characterization of PD-GSC response to pitavastatin. UMAP plots of scRNA-seq profiles, annotated according to treatment conditions (untreated control, vehicle – DMSO, and pitavastatin – PSTAT), for **(A)** SN520 and **(B)** SN503. Scatter plots show proportions of each subtype in each PD-GSC population across treatment for **(C)** SN520 and **(D)** SN503. **(E-F)** Flow cytometry analysis of PN and MES markers CD133 (PN) and CD44 (MES) across pitavastatin-treated cells for SN520 and SN503, respectively. Proportions of cells positive for each subtype marker are quantified in the adjacent barplots underneath. **(G-H)** Heatmap of inferCNV scores for SN520 and SN503, respectively. Cells (rows) are grouped based on treatment conditions (same color annotation as in (A) and (B)). Genes (columns) are arranged according to their chromosomal positions. **(I)** Dose-response curves of naïve SN520 PD-GSCs (light blue) and SN520 PD-GSCs that survived an initial pitavastatin-treatment (treated – dark blue). Adjacent plot shows corresponding AUC values from dose-response curves generated from subsequent PD-GSC cultures derived from original pitavastatin– or vehicle-control-treatment for SN520 (left) and SN503 (right). Paired t-test results showed a sustained (significant) increase in AUC values of the PSTAT-treated SN520 PD-GSCs relative to their vehicle-control counterparts but not for SN503.

Interestingly, GSVA enrichment scoring (Supplementary Figure 3) showed that while the relative proportions of cells for each molecular subtype (i.e., CL, PN, MES) was fairly consistent in vehicle control, the 4-day pitavastatin treatment of SN520 responder cells showed a dramatic increase in the proportion of cells of the MES subtype (Figure 2C). In stark contrast, the subtype composition of the SN503 non-responder cells remained relatively constant during treatment with pitavastatin and vehicle control (Figure 2D). Notably, the longitudinal patterns of subtype composition within each PD-GSC population determined from scRNA-seq time course analysis were inconsistent with findings from bulk-RNA-seq analysis. Cytometry analysis confirmed findings from scRNA-seq analysis that pitavastatin treatment of SN520 resulted in an increase in the proportion of CD44+ (MES) cells from 28.2% to 65.35%, and a simultaneous decrease in CD133+ (PN) cells from 52.7% to ∼1%. Of note, SN520 had a sizeable (35.3%) proportion of CD133+/CD44-PN cells, which were nearly eliminated by D4 (Figure 2E), likely due to a combination of treatment-induced killing and a transition of surviving cells to a MES state. By contrast, pitavastatin treatment did not cause a change in the proportion of CD44+ cells in SN503 (87% on D1 to 85.11% on D4, Figure 2F). The significant decrease in the relative proportion of CD133+ cells within SN503 (from 38.1% on D1 to 9.51% on D4), especially over the first two days of treatment, was likely due to pitavastatin-induced killing of a susceptible PN subpopulation (*9*). Interestingly, the relative proportion of CD133+/CD44-PN cells (1.41%) within SN503 was negligible; pitavastatin sensitivity appeared to be associated with a CD133+/CD44+ sub-population that was in higher abundance (36.7%).

To differentiate between selection and differential proliferation as the mechanism responsible for the observed shifts in subtype composition, we used canonical cell cycle gene expression signatures to score each cell (METHODS) and found that only small proportions of cells within each PD-GSC culture were in the S or G2/M phase regardless of treatment context (Supplementary Figure 4). Consistent with this finding, cytometry-based DNA quantification of individual cells confirmed that only a small proportion of cells across both PD-GSCs were actively proliferating during pitavastatin treatment (Supplementary Figure S5). Theoretical calculations based on cell division rate and treatment duration (Supplementary Figure S6), as well as the homogeneity of CNV states pre– and post-treatment of both PD-GSCs (Figure 2G, H) both independently suggested that cell subtype transitions of surviving SN520 cells, rather than a natural selection and expansion, was responsible for the observed treatment-induced changes in subtype composition and phenotypic characteristics. Finally, overall drug sensitivity of surviving SN503 cells remained relatively unchanged post-pitavastatin treatment for ∼30 days (Figure 2I; paired t-test p-value = 0.348). In stark contrast, there was significant log2-fold increase of 2.42 in IC_50_ of surviving SN520 cells from 0.42 μM to 2.24 μM, which was sustained over 100 days (Figure 2I paired t-test p value = 1.526e-05), demonstrating the long-term functional consequences of drug-induced PMT.

### Characterization of transcriptional states of PD-GSCs reveals multiple mechanisms of primary and acquired resistance

Dimensionality reduction with PCA and subsequent Louvain clustering (METHODS) of differentially expressed genes (DEGs, Supplementary Figure S7) organized the 5,402 SN520 cells into 14 clusters (Figure 3A, B) and the 5,722 SN503 cells into 12 clusters (cl_503/520_-*i*; Figure 3C, D). As expected, the SN520 Louvain clusters were predominantly comprised of either vehicle– or pitavastatin-treated PD-GSCs (Figure 3E). By contrast, several SN503 Louvain clusters contained a mix of both vehicle– and drug-treated cells (Figure 3F). Below we summarize findings based on pathway enrichment analysis of DEGs within each Louvain cluster (Figure 3G). A more detailed description is included in the Supplement.

**Figure 3.**
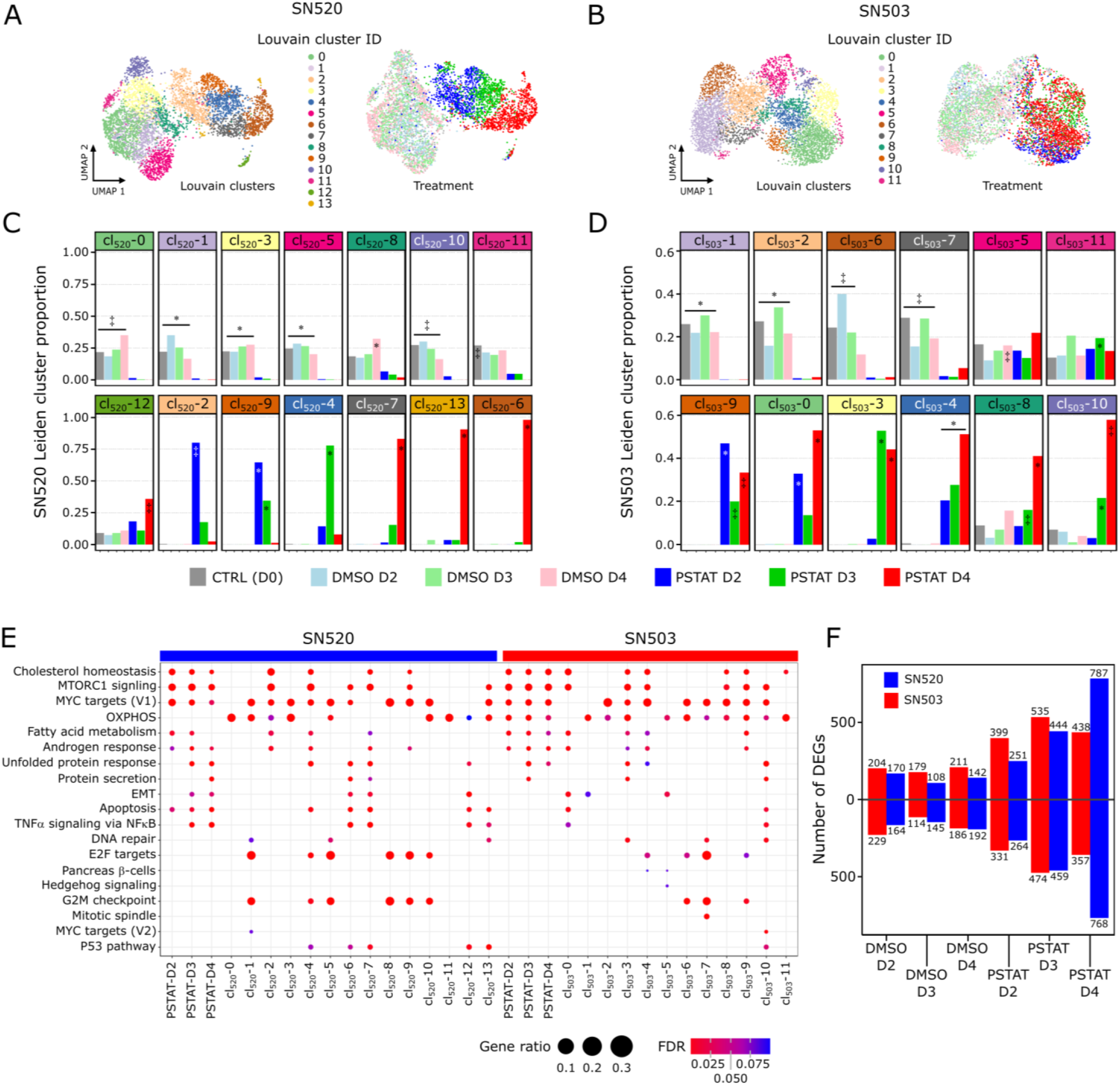
Differential expression and pathway enrichment analysis reveals underlying processes driving pitavastatin responses. (**A**) Heatmap of the top upregulated DEGs, based on FDR p-values, across the 14 Louvain cell clusters (cl) identified in vehicle-control– and pitavastatin-treated SN520 PD-GSCs. Adjacent UMAP plot with treatment annotation (same as Fig 2A) included for reference. **(B)** Corresponding UMAP plots of scRNA-seq profiles annotated according to Louvain cell cluster (left) and treatment condition (right) as reference. **(C)** Cell proportions for each Louvain cluster that belong to each treatment condition for SN520. Significant enrichment of treatment condition within Louvain cluster indicated by asterisk (FDR ≤ 0.05) or double dagger (FDR ≤ 1e-05) **(D)** Cell proportions for each Louvain cluster that belong to each treatment condition for SN503. Significant enrichment notation identical to that used in (D). **(E)** Dotplot of hallmark gene sets enriched across SN503 and SN520 PD-GSCs, grouped with respect to either drug-treatment duration or Louvain clustering. Dot size represents the ratio of number of upregulated genes associated with a PD-GSC grouping to the number of genes associated with a specific hallmark gene set. Dot colors indicate significance of enrichment (FDR value). **(F)** Total number of up– and down-regulated DEGs, relative to untreated control (D0) cells, at each treatment time point for SN503 (red) and SN520 (blue).

*SN520 Clustering & Enrichment*. Consistent with the mechanism of action of pitavastatin, gene set enrichment analysis (GSEA, Supplementary Tables S1-S2) revealed that within two days upon initiation of treatment SN520 cells differentially regulated cholesterol homeostasis, biosynthesis, and maintenance, as well as MTORC1 signaling. Day 3 onwards the cells differentially regulated stress response genes including unfolded protein response, protein secretion, P53 pathway, and apoptosis. Interestingly, upregulation of both apoptosis and EMT genes across subpopulations of drug-treated D4 cells (cl_520_-6, cl_520_-7) was consistent with simultaneous induction of these pathways by TGFβ during tumor formation and progression, with cell fate being dependent on cell-cycle phase (*35*, *36*). In this case, cl_520_-6 and cl_520_-7 cells were in G1/S phase, suggesting that SN520 cells escaped apoptosis by transitioning into the MES subtype (Supplementary Figure S7).

*SN503 Clustering & Enrichment*. Although there were fewer DEGs in SN503 as compared to SN520 (Figure 3H), the Louvain clusters of pitavastatin-treated SN503 cells did bear similarity to SN520 clusters with regard to differential regulation of certain pathways, including cholesterol homeostasis, fatty acid metabolism, MTORC1 signaling, androgen response, and unfolded protein response (Supplementary Tables S3-S4). However, the differential expression patterns were distinct between the two PD-GSCs. For instance, pitavastatin-treated SN503 cells did not cluster by treatment time, instead cells from all time points grouped together across multiple Louvain clusters (Figure 3C, F) characterized by upregulation of oxidative phosphorylation (OXPHOS, Figure 3G, Supplementary Table S3), which has been associated with drug resistance in tumor cells (*37–40*). Moreover, only a small proportion of pitavastatin-treated SN503 cells differentially regulated EMT-associated genes (cl_503_-0 and cl_503_-5) (Figures 2, 3H). These findings suggested that different regulatory mechanisms were likely responsible for the distinct differential expression patterns of key pathways, as well as the responder and non-responder phenotypes of SN520 and SN503, respectively.

### Inference and dynamic simulation of transcriptional regulatory networks identifies mechanisms driving cell-state changes and intervention strategies

We applied single-cell SYstems Genetics Network AnaLysis (scSYGNAL) framework to uncover the transcriptional regulatory networks (TRNs, (*41*, *42*)) responsible for driving the distinct transcriptome responses of the two PD-GSCs. Briefly, Mechanistic Inference of Node Edge Relationships (MINER), an algorithm within the scSYGNAL framework, was used to identify modules of genes (regulons) that were co-regulated differentially in response to treatment across sub-populations of cells (*43*, *44*). Further, using the transcription factor binding site database (*45*) and the Framework for Inference of Regulation by miRNAs (FIRM, (*46*)), scSYGNAL implicated specific TFs and miRNAs in mechanistically co-regulating genes of all regulons. Post-processing of the resulting TRNs using MINER (*47*) clustered regulons with similar activity profiles across subpopulations of cells into transcriptional programs ( Pr_503/520_-*i*) and clustered single cells with similar program activity profiles into distinct transcriptional states (St_503/520_-*i*). Here onwards we will refer to the TRNs for each PD-GSC as scSYGNAL-520 and scSYGNAL-503.

*scSYGNAL-520* modeled the influence of 109 TFs and 505 miRNAs in mechanistically regulating 1,668 genes across 572 regulons that organized into 19 transcriptional programs and were differentially active across 17 transcriptional states (Fig. 4A; Supplementary Table S5-S6). Strikingly, nearly every transcriptional program was enriched for genes that have been shown to be essential to GSC survival (Supplementary Table S7, (*48*)). GSEA revealed that many pathways identified within Louvain clusters were recapitulated by programs (Figure 3G, Supplementary Table S8). For instance, Program 0 (Pr_520_-0) – the largest program consisting of 169 regulons, was enriched for genes associated with cellular stress responses, including unfolded protein response, androgen response, p53 pathway, and apoptosis. Pr_520_-1, the second largest program (61 regulons) was enriched for cholesterol homeostasis and MTORC1 signaling. Pr_520_-2 (proliferation), Pr_520_-5 and Pr_520_-6 (TNFα signaling via NFκB) showed variable activity in states enriched with vehicle-treated cells, but were uniformly underactive in states enriched with pitavastatin-treated cells (Figure 4A). Only four states (St_520_-0 – St_520_-3) were enriched for D3 and D4 pitavastatin-treated cells (Figure 4B), suggesting that they might represent drug resistant states adopted by the surviving subpopulation of cells to avoid pitavastatin-induced killing. Furthermore, when transcriptional states were rearranged with respect to their predominant treatment condition, program activities increased (nearly) monotonically over the course of treatment, which suggested that treatment-induced state transitions occurred through continuous rather than discrete changes in expression in SN520 (Figure 4C, Supplementary Figure S8).

**Figure 4.**
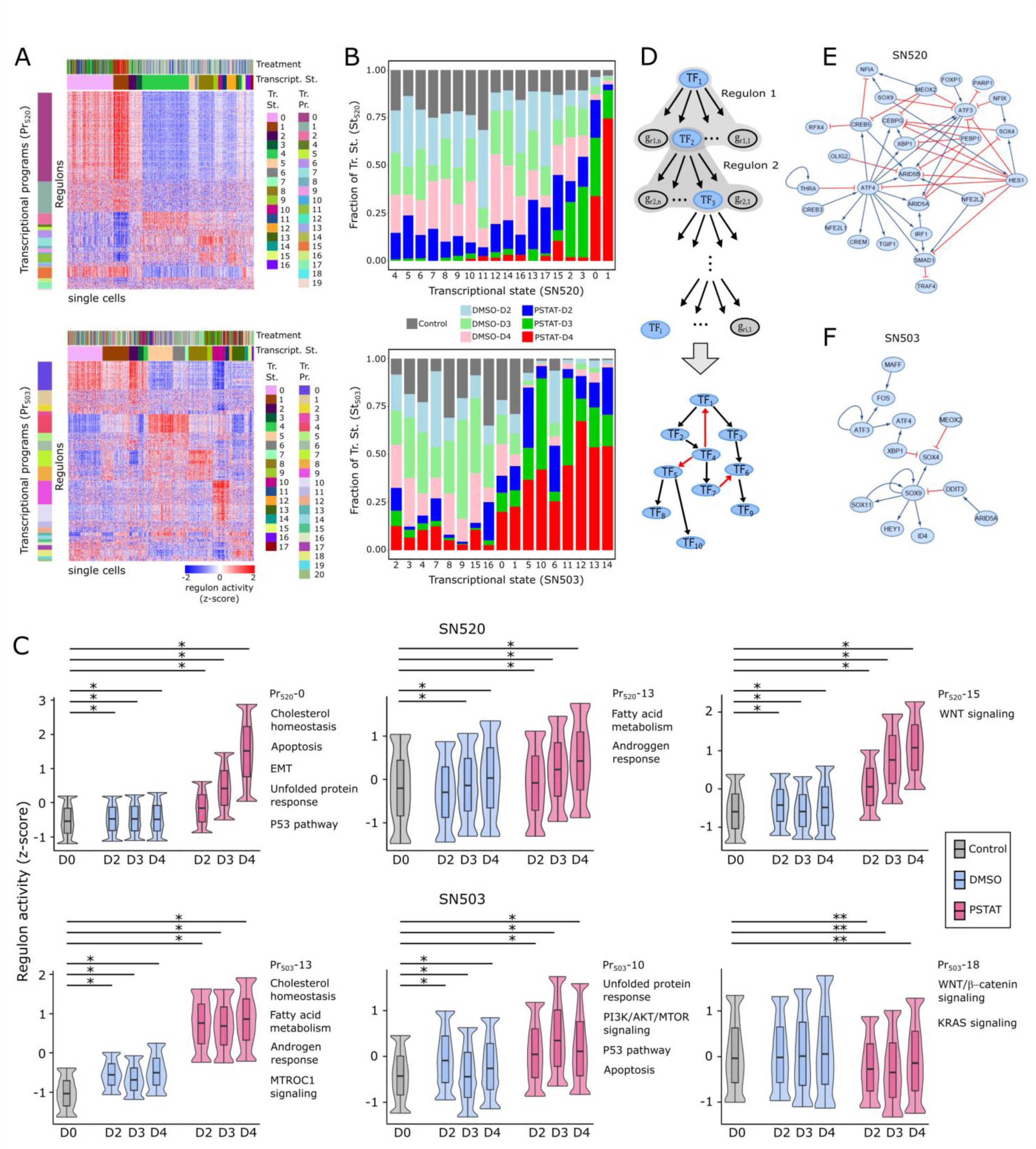
MINER3 transcriptional regulatory network inference reveals mechanisms of cell-state changes. **(A)** Heatmaps of normalized regulon activities across SN520 (top) and SN503 (bottom) PD-GSCs. Regulons (rows) are organized into transcriptional programs (Pr) while single cells (columns) are organized into transcriptional states (St). Left-adjacent color bars indicate what regulons belong to a particular transcriptional program. Left-adjacent color bar indicates transcriptional programs. Top color bars indicate treatment condition (color annotation identical to Fig. 1E) and corresponding transcriptional state for a single cell. (**B)** Stacked barplot show proportion of cells within each transcriptional state from each treatment condition for SN520 (top) and SN503 (bottom). **(C)** Boxplot/violin plots of distributions of regulon activity for select programs across treatment conditions for SN520 and SN503. Regulon activity values were capped between the lower 2.5% and 97.5% range of values. Labels indicate program IDs and select hallmark gene sets (*90*) enriched within each program. The box represents the inter-quantile range (IQR – 25^th^ and 75^th^ percentile) and median activity value while the whiskers represent 1.5x IQR. Asterisks indicate statistically significant differences between regulon activity distributions. Single asterisks (*) denote activity distribution of untreated controls (CTRL) is significantly lower than distribution being compared (FDR << 1e-3). Double asterisks (**) denote distribution of untreated controls is significantly higher than either vehicle-treated (DMSO) or pitavstatin-treated (PSTAT) distributions being compared (FDR << 1e-3). **(D)** Flow diagram outlining approach to derive core TF-TF network from MINER3 results. Final core TF-TF networks derived for **(E)** SN520 and **(F)** SN503.

*scSYGNAL-503* modeled the regulation of 1,875 genes by 114 TFs and 507 miRNAs across 420 regulons, organized into 21 distinct transcriptional programs, whose activity profiles stratified SN503 cells into 17 transcriptional states (Figure 4A bottom heatmap, Supplementary Tables, S9-S10). Like SN520, a large portion of these programs were enriched with essential genes for GSC survival (Supplementary Table S11; (*48*). Several programs were similar to those identified in SN520, including Pr_503_-13 (cholesterol homeostasis, MTORC1 signaling and fatty acid metabolism), Pr_503_-9 and Pr_503_-10 (stress responses, including vesicle-mediated transport, unfolded protein response, and p53 pathway). In contrast to SN520, many SN503 programs were uniquely enriched in distinct processes, including WNT/β-catenin and KRAS signaling (Pr_503_-18, Fig. 4F, Supplementary Table S12). Unlike SN520, D3 and D4 pitavastatin-treated SN503 cells co-clustered in significant proportions with untreated and vehicle-treated cells across >75% of the 17 states, suggesting that a large number of SN503 cells may have been in pitavastatin-resistant states even prior to drug exposure (Figure 4C). Interestingly, multiple states included pitavastatin-treated cells from all time points, including seven states in which the drug-treated cells represented >50% of all cells (Figure 4B). The seven transcriptional states were distinct in their activity patterns of some programs, including Pr_503_-4 (apoptosis, EMT, IL6/JAK/STAT3 signaling), which was overactive in St_503_-5, St_503_-6, and St_503_-10; and Pr_503_-10 (MTORC1 signaling, hypoxia, and unfolded protein response), which was overactive in St_503_-10 and St_503_-11. The heterogeneous activity patterns of these programs, which were enriched for processes linked to chemotherapeutic resistance (*49*), suggests that multiple mechanisms likely contributed to pitavastatin resistance in SN503.

#### Core TF-TF interaction networks governing PD-GSC response to pitavastatin

We derived a “core” network of TF interactions to investigate how transcriptional regulatory mechanisms contributed to PMT and pitavastatin resistance (Figure 4D). Each directed TF-TF interaction was categorized as activating or repressing based on positive or negative pairwise correlation of expression levels between two TFs, respectively. The topology of the core TF network for each PD-GSC population was distinct (METHODS), with 56 interactions (edges) among 31 TFs (nodes) in scSYGNAL-520 and only 13 interactions connecting 15 TFs in scSYGNAL-503 (Figure 4E, F). Multiple TFs in the core scSYGNAL-520 TF network have been linked to response-relevant processes including EMT, cell differentiation, adaptive responses, and stem-cell maintenance (Supplementary Table S13). Nine TFs were common between the core networks (overlap p-value: 9.44e-05), including ARID5A, ATF3/4, MEOX2, SOX9, XBP1, and HEY1, a Notch signaling regulator. TFs unique to the core scSYGNAL-503 network included DDIT3, MAFF, STAT3, and ID4, which have been implicated in multiple GBM-relevant processes, (Supplementary Table S13). Notably among these TFs, ID4 has also been shown to play a role in the pathogenesis of GBM, driving tumor-initiating cell formation by increasing two key cell-cycle and differentiation regulatory molecules – cyclin E and Jagged 1 (*50*). These findings suggest that the core networks captured TF-regulation that play central roles in GBM and gliomas in general.

### Trajectory analysis and network simulations uncover mechanisms of primary and acquired resistance

Using Monocle3 we discovered that pseudotemporal ordering of SN520 cells correlated with treatment duration and concomitant drug-induced PMT (Pearson correlation coefficient *r* = 0.723). We observed similar agreement between treatment duration and inferred trajectories from RNA velocity analysis (*51*), as velocity vectors pointed towards 4-day treated cells (Figure 5A). In parallel, we calculated the critical transition index (*I_c_*), a quantitative metric of the high-dimensional state of a system that predicts whether a cell population is undergoing a state transition (higher *I_c_* values) or if it has reached some stable attractor state (lower *I_c_* values) (*52*). *I_c_* values of SN520 decreased during drug treatment but remained relatively constant in the vehicle control (Figure 5B), indicating that pitavastatin had driven the entire PD-GSC population into a predominantly drug-resistant MES subtype attractor state. By contrast, pseudotemporal ordering of SN503 cells did not correlate with treatment time (Pearson correlation coefficient *r* = –0.0167,) and was associated with high *I_c_* values throughout the course of the experiment for both vehicle control and drug treatment, likely driven by the higher heterogeneity of the cells. Consistently, these GSCs exhibited a rather turbulent vector field where RNA velocities projected into multiple directions (Figure 5A). Modeling concerns associated with pseudotime and trajectory inference analysis notwithstanding, e.g., hyperparameter optimization (*53*, *54*), the pseudotime and criticality analyses demonstrated stark contrast between the responses of the two PD-GSCs; SN520 exhibited concerted pitavastatin-induced state transitions, relaxing into a regulated state, while SN503 exhibited a seemingly disorganized response without concerted transition of all cells into an attractor state.

**Figure 5.**
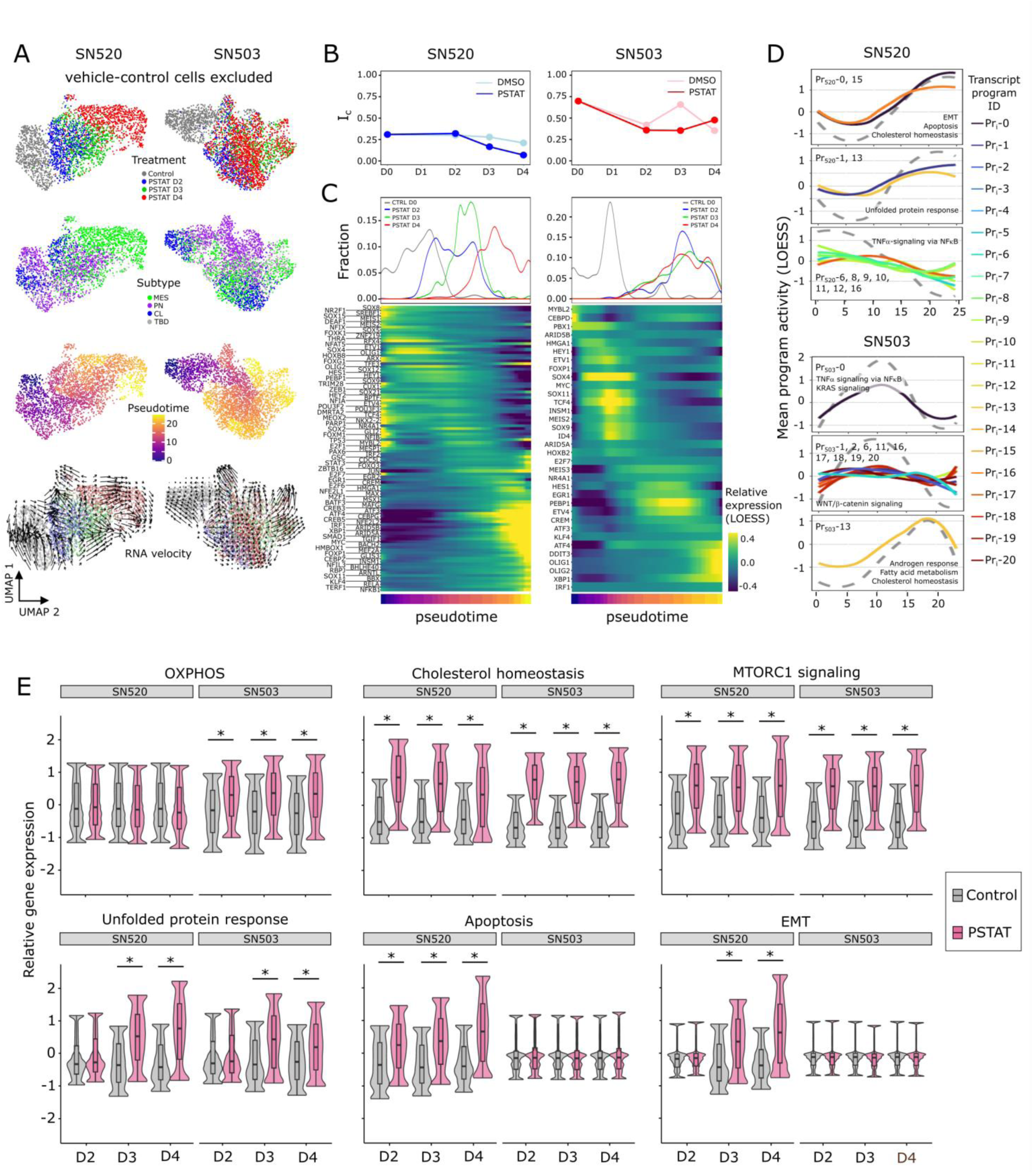
Distinct trajectories define SN520 and SN503 pitavastatin response. (**A**) UMAP plots of vehicle– and pitavastatin-treated cells for SN520 (left column) and SN503 (right column). Annotation highlight treatment conditions (top row), molecular subtype (2^nd^ row), pseudotime (3^rd^ row) and RNA velocity (4^th^ row). **(B)** Critical transition index (I_c_) of SN520 (blue) and SN503 (red) cells treated with vehicle (DMSO – light) or pitavastatin (PSTAT – dark). **(C)** LOESS regression of TF expression behavior sorted according to peak expression along pseudotime. Density plots depict distribution of sample time points along pseudotime trajectory. Heatmap shows expression of TFs rank sorted by time of peak expression along pseudotime (color bar beneath heatmap). **(D)** Select set of LOESS regression of mean program activities with respect to pseudotime. Regulons are clustered based on their dynamic activity profiles with respect to pseudotime. Dashed grey line represents the average shape of the curves for each cluster. Labels indicate which transcriptional programs were grouped into each cluster. Select hallmark gene sets (*90*) enriched within programs are labeled as well. **(E)** Boxplots/violin plots of expression of genes associated with indicated pathways/processes (*90*) on respective treatment days. Relative gene expression values were capped at the lower 2.5% and 97.5% range of values. Labels indicate select hallmark gene sets enriched within subpopulation of cells (treatment time point). Asterisks indicate statistically greater expression in pitavstatin-treated cells (PSTAT) relative to untreated control (CTRL) counterparts (Wilcoxon rank test, FDR << 1e-5). The box represents the inter-quantile range (IQR – 25^th^ and 75^th^ percentile), median activity value while the whiskers highlight 1.5x IQR.

To identify putative drivers of treatment response, we performed LOESS regression and rank ordered TFs with respect to timing of peak expression along the pseudotime trajectories and uncovered a distinct sequence of changes in the activity of multiple TFs in each PD-GSC population (Figure 5C). Within SN520, multiple TFs previously associated with PMT in GBM (e.g., ATF3, CREB, and NFE2L2) positively correlated with pseudotime trajectory (Supplementary Table S13 – Moran’s I value). Notably, the rank order of TFs in SN520 was quite different from previously proposed sequence of transcriptional events driving PMT (*55*), which highlights the diversity of regulatory mechanisms that have been implicated in driving EMT in multiple cancers (*56*, *57*). As expected, we did not observe temporal sequence of changes in expression levels of TFs across SN503 cells (Figure 5C, Supplementary Table S13).

In addition, we investigated the consequence of differential expression patterns of TFs by examining, along pseudotime trajectories, the dynamic activity patterns of transcriptional programs that they regulated (Figure 5D, Supplementary Figure S9). Activity of the stress-response-associated programs (Pr_520_-0) increased along the pseudotime trajectory of SN520 cells, implicating 80 associated TFs, including ATF3, ATF4, CREB3, CREB5, JUN, KLF4, MYC, SOX4/9, and TCF4. In the case of SN503, we identified multiple treatment-activated programs for key processes (Figure 4C) including unfolded protein response and OXPHOS (Pr_503_-9 and Pr_503_-10), cholesterol regulation (Pr_503_-4) and EMT (Pr_503_-5 and Pr_503_-13) that showed upregulated gene expression relative to the untreated control condition (Figure 5E). Importantly, scSYGNAL-503 had accurately identified TFs that have been mechanistically implicated in regulation of these processes, such as AR, FOS, MYC, TP53, and E2F7 for Pr_503_-9 and Pr_503_-10 (*58*).

#### Ensemble modeling and analysis of GSC states via simulated TF-TF network dynamics

We performed *in silico* perturbations on the core TF-TF networks using the random circuit perturbation (RACIPE) algorithm (*59–61*) to identify transcriptional regulatory mechanisms that governed pitavastatin-induced cell state changes across the two PD-GSCs (Figure 4D, E). RACIPE was originally developed to investigate EMT circuits in cell development and other cancers by creating an ensemble of dynamic models based on ordinary differential equations and Hill function kinetics (*62–64*). First, we tested whether the TF-TF network model for each PD-GSC could accurately predict their observed pitavastatin-induced cell states using untreated (D0) TF expression levels to initialize the network. By performing 1,000 RACIPE simulations, we determined that the simulated stable steady states were statistically similar to the observed cell states of each PD-GSC on D4 of pitavastatin treatment (Figure 6A, B, Supplementary Figure S10).

**Figure 6.**
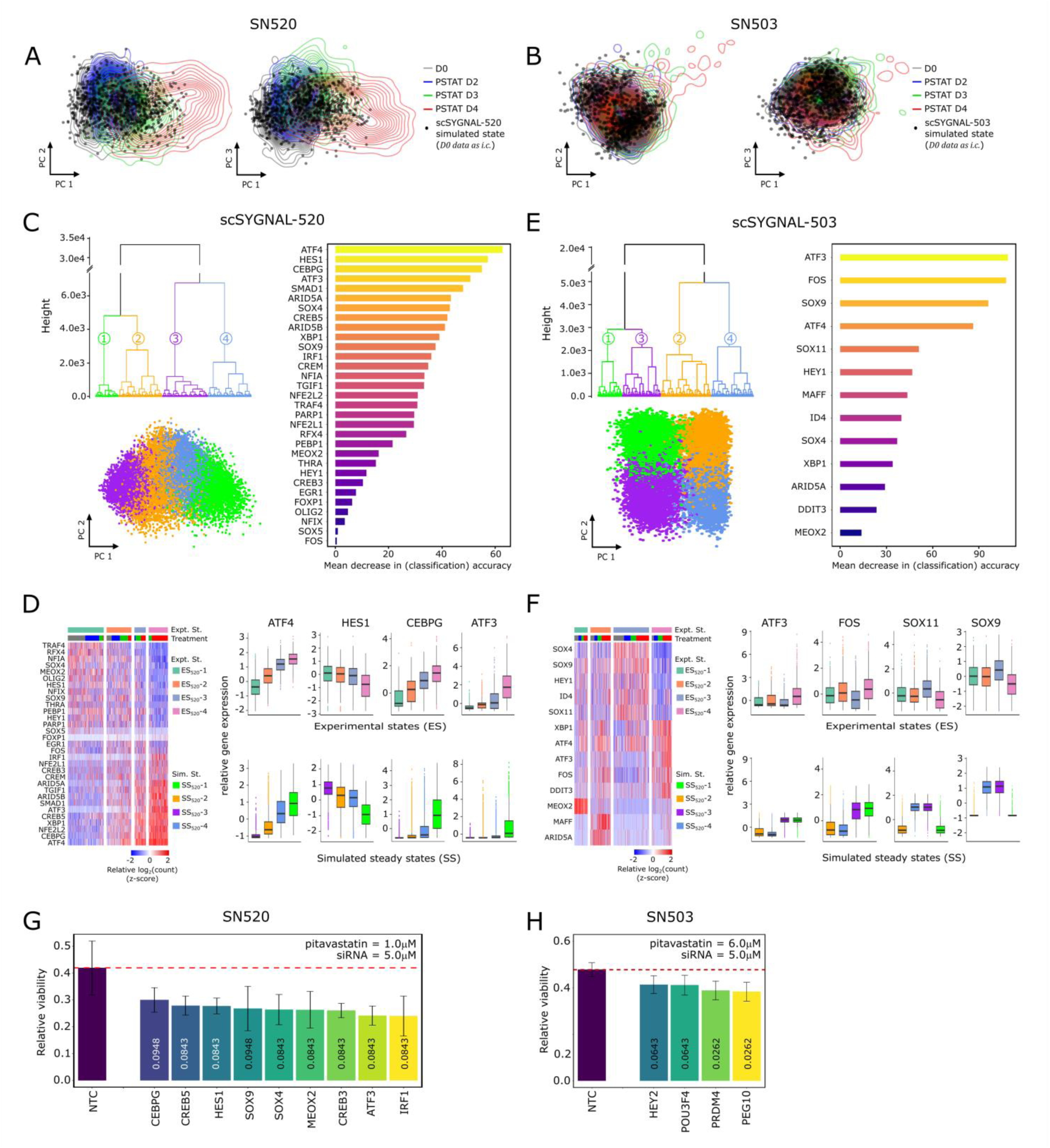
Dynamic simulations of core TF regulatory network supports phenotypic plasticity of GSCs. Simulated transcriptional states projected along first two principal components. Contour lines represent distribution of scores from PCA of TF expression states from single PD-GSCs for **(A)** SN520 and **(B)** SN503. One thousand simulated states were generated using scSYGNAL-520/503 as network topologies and using respective D0 scRNA-seq data as inputs to RACIPE algorithm. **(C)** Three plots summarize results from 1 million simulations using scSYGNAL-520 and randomly selected initial conditions as inputs to RACIPE algorithm to explore plausible steady states supported by network topology derived from MINER3 (simulations are distinct from those in (A)). Dendrogram highlights four distinct simulated steady states generated by RACIPE using core TF network and randomly selected initial conditions as input.. Simulated states projected along first two PCs. Horizontal barplot visualizes rank-ordered importance of TFs in distinguishing four simulated states per random forest analysis based on the mean decrease in accuracy in categorizing sample if the TF were excluded from the model. **(D)** Heatmap of expression for TFs that define core TF network in SN520 cells that define experimental states (ES_520_-i) used as basis of comparison for simulated states. Adjacent boxplots of top four most important TFs per random forest analysis. Top row of boxplots show distributions of expression of TFs for each experimental states identified. Bottom row includes distributions of simulated expression values (normalized) from simulations that used randomized initial conditions. **(E-F)** Same as (C-D), but for SN503. **(G)** Viability of SN520 following 4-day treatment with 1.0 μM pitavastatin and simultaneous siRNA-mediated KD of TFs or non-template control (NTC – red dashed line). Values in bars represent FDR p-values indicating significant decrease relative to NTC. Values in bars represent FDR p-values indicating significant decrease relative to NTC siRNA condition. Adjacent scatterplot compares rank ordering of tested TFs to corresponding rank ordering based on predicted change in proportion of states belonging to MES state 1. TFs below the diagonal have a greater impact on reducing viability than was predicted. **(H)** Viability of SN503 following pitavastatin treatment and simultaneous siRNA-mediate KD of TFs associated with OXPHOS-associated relative to NTC and 4-day treatment 6.0 μM pitavastatin treatment. Again, values in bars represent FDR p-values. Values in bars represent FDR p-values.

We then investigated how the core TF network contributed to phenotypic plasticity by determining the range of steady states that could emerge from each network topology. We simulated 10,000 distinct models (i.e., parameter sets) across 100 randomly selected initial conditions resulting in an ensemble of 1 million simulations for each PD-GSC population, which was sufficient to yield convergent solutions (Supplementary Figure S11 (*59–61*)). Based on pairwise Euclidean distances (METHODS) and hierarchical clustering, all simulated states generated by the core TF network for SN520 clustered into four distinct steady states (Figure 6C). The simulated states stratified along the first principal component, recapitulating a continuum of progression from a PN to MES state (Figure 6C). Pairwise comparisons of mean expression profiles of the core network TFs demonstrated that the simulated states were statistically similar to experimentally observed PD-GSC states (Figure 6C, Supplementary Figure S10). Supervised classification using random forest analysis further revealed that ATF3/4, CEBPG, and HES1 contributed the most to distinguishing the four simulated states (Figure 6C), which mirrored expression behavior across experimental data for SN520 (Figure 6D).

RACIPE simulations for SN503 also yielded four distinct stable steady states that did not show a gradient in PCA space as in the case of the SN520 simulated states (Figure 6E). Three of these states were similar to two experimentally observed PD-GSC states (Figure 6E) associated with elevated expression of SOX4, SOX9, SOX11, HEY1, and ID4 (simulated states 3 and 4 and experimental state 4), or elevated expression of ATF4, ATF3, and FOS (simulated states 1 and 3 and experimental state 4). The experimentally observed states not identified by RACIPE simulations were associated with elevated expression of MEOX2, MAFF, and ARID5A, which were “root” nodes, i.e., TFs without any upstream regulators in the context of the model. Consequently, expression of these TFs in the RACIPE simulations was dependent upon the randomly selected initial conditions. However, the subset of simulations in which MEOX2, MAFF, and ARID5A had elevated initial conditions generated states that were indeed similar to experimentally observed states ES_503_-1 and ES_503_-2 (Supplementary Figure S10). Further, for distinguishing the four SN503 PD-GSC states, random forest analysis identified MEOX2, MAFF, and ARID5A as the most important TFs, followed by ATF3, SOX9, and SOX11 (Supplementary Figure S10). Interestingly, all of these TFs have previously been implicated in tumor stemness, progression, invasiveness or resistance, suggesting multiple mechanisms may have contributed to pitavastatin resistance in SN503 (Supplementary Table S13).

#### In silico network perturbations implicate specific TFs in mechanistically driving treatment-induced cell state transitions and drug resistance in PD-GSCs

After benchmarking the random forest models as 85% and 90% accurate in predicting cell states of SN520 and SN503, respectively (Supplementary Figure S12), we used them in perturbation simulations to identify mechanistic drivers of treatment response of each PD-GSC. Specifically, we performed an additional 1 million RACIPE simulations to model the consequence of 95% knockdown in each TF within the core network on treatment-induced change in the relative abundance of each of the four steady states for the two PD-GSCs. (Supplementary Figure S13). This analysis predicted that knockdowns in each of ten TFs, viz., ATF4, IRF1, NFE2L2, CREB3, XBP1, ARID5A, SMAD1, CREB5, CEBPG, and ATF3, would result in significant reduction in the relative abundance of simulated states with large subpopulations of MES subtype cells in SN520 (Figure 6G). Notably, all ten TFs have been implicated in driving EMT across different cancers, including GBM (Supplementary Table S13). RACIPE simulations predicted that decrease in the proportion of MES subtype-associated cell states in SN503 was likely through perturbations in just two TFs, namely SOX9 and SOX11 (Supplementary Figure 13) both of which were also implicated in driving PMT (Supplementary Table S13).

#### siRNA knockdowns of TFs validate core TF networks

We tested RACIPE predictions by investigating whether siRNA (Dharmacon^TM^) knockdown of TFs during pitavastatin treatment would block PMT leading to synergistic increase in PD-GSC killing. Indeed, knockdowns in nine TFs (5/10 predicted), including ATF3, IRF1, CREB3, CREB5, and CEBPG, significantly potentiated pitavastatin killing of SN520 (Figure 6I). Notably, increased cell death of SN520 was observed only when siRNA and pitavastatin were administered simultaneously, but not when cells were pre-treated with siRNA prior to pitavastatin treatment (data not shown). Given that siRNA knockdown is typically manifest in protein reduction maximally in 2-3 days post-transfection, dynamic induction of TF activity by pitavastatin appears to have been essential for achieving the TF knockdown effect on SN520 PD-GSC survival. In stark contrast, none of the TF knockdowns had any consequence on viability of SN503 (Figure 6H, J). Altogether, the experimental findings corroborated the roles of nine TFs implicated by scSYGNAL and RACIPE analysis in driving PMT, thereby conferring pitavastatin resistance in SN520, but not in SN503, wherein a large fraction of the cell population was in a drug resistant MES state, even prior to drug treatment. As an alternative approach, we identified 24 additional TFs by MINER as important for mechanistically upregulating putative resistance mechanisms, including OXPHOS (Figure 2G, Supplementary Table S3, S12), and discovered that knocking down four TFs (HEY2, POU3F4, PRDM4, and PEG10) indeed potentiated pitavastatin killing of SN503, likely by disrupting one or more primary resistance mechanism(s) (Figure 6K).

### Trajectories towards acquired resistance expose vulnerabilities to secondary drugs

Finally, we investigated whether knowledge of mechanistic drivers of PMT could enable rational selection of a second drug that could potentiate the action of pitavastatin. Using Open Targets (*65*), we identified eight drugs that targeted TFs and genes associated with pitavastatin-induced PMT trajectories in SN520. We hypothesized that pitavastatin-induced cell state changes place cells in transitional states that may expose new vulnerabilities that could be targeted by secondary drugs. We selected vinflunine, a vinca alkaloid that binds to tubulin and inhibits microtubule polymerization, thereby inducing G2/M arrest and ultimately apoptosis. Originally developed to treat advanced or metastatic transitional cell carcinoma of the urothelial tract (*66*), vinflunine has been tested in multiple Phase III trials for many cancers, used as a likely potentiator of anti-cancer effects of other drugs (*67*). Based on vinflunine’s mechanism of action, we identified multiple regulons containing tubulin-related genes (for example, SN520 regulons R_520_-0 and R_520_-43; SN503 regulons R_503_-19, R_503_-38, and R_503_-52). In SN520, the activity for R_520_-0 and R_520_-43 increased significantly in response to pitavastatin (Figure 7A). By contrast, pitavastatin-induced upregulation of tubulin-associated regulons was varied across in SN503, with only R_503_-19 showing consistent over activity across all time points. R_503_-38 showed significantly higher activity in pitavastatin-treated cells relative to vehicle-treated, with maximal activity on D3. Finally, R_503_-52 activity levels were slightly higher relative to vehicle control (Figure 7B). The ability of vinflunine to block pitavastatin-induced cell state transitions was investigated in three experimental designs, one in which both drugs were added simultaneously and the other two in which vinflunine was added at 24 or 48 hrs after initiation of pitavastatin treatment to match the timing when pitavastatin-treatment induced the highest activity of tubulin regulons (Figure 7C). The efficacy of the drug combinations were compared to outcome of treatments of PD-GSCs with each individual drug.

**Figure 7.**
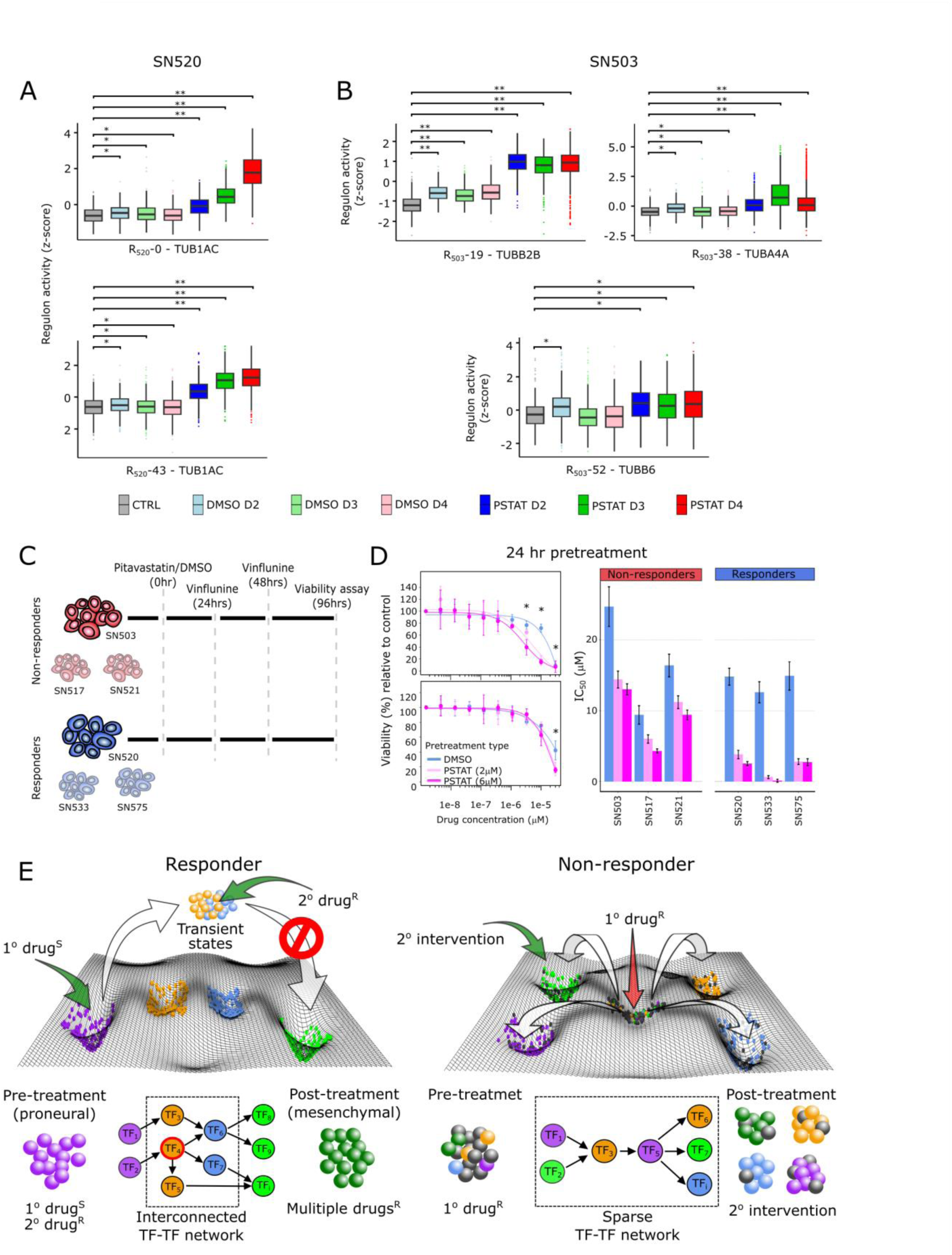
Dynamic behavior of regulons reveal additional targets that guide rational secondary drug selection. (**A**) Distribution of activities for representative tubulin-associated regulons across SN520 PD-GSCs. Statistically significant differences in regulon activity, relative to activity in untreated control cells are indicated by asterisks or double daggers (Wilcoxon rank test, * FDR ≤ 1e-20, ** FDR ≤ 1e-150). **(B)** Distribution of activities for representative tubulin-associated regulons across SN503 PD-GSCs across treatment conditions. Asterisks indicate conditions having significantly higher values relative to untreated controls (Wilcoxon rank test, * FDR ≤ 1e-20, ** FDR ≤ 1e-150). **(C)** Experimental designs used to test effects of sequential pitavastatin and vinflunine treatment on non-responder and responder PD-GSC populations. **(D)** Dose-response curves for SN520 (top) and SN503 (bottom) resulting from a 24hr pre-treatment with vehicle or pitavastatin (2μM or 6μM) followed by vinflunine treatment (1.5e-9, 4.6e-9, 13.7e-9, 41.2e-9, 123.5e-9, 370.4e-9, 1.10e-6, 3.30e-6, 10.0e-6 30.0e-6 M). Adjacent barplot of IC_50_ values determined from 24hr pretreatment with vehicle or pitavastatin (2μM or 6μM) for all non-responder and responder PD-GSCs tested. Results from 48hr pretreatment are included as Supplementary Figure S14. **(E)** Depiction of how topologies of the underlying response TF-TF networks, in response to drug treatment, can drive responder and non-responder PD-GSCs to transition into different states along a Waddington-like phenotypic landscape. Treatment with a drug to which cells are sensitive (1° drug^S^) activates a highly interconnected network of a responder PD-GSC, driving PMT across a majority of the surviving cell population, enabling acquisition of resistance to “multiple drugs^R^”. Secondary intervention with a drug to which cells are resistant (2° drug^R^) to target vulnerabilities in the intermediate states potentiates killing and likely blocks PMT. By contrast, the non-responder PD-GSC is comprised of sub-populations of cells that are already in states (center well) that are resistant to the primary drug (1° drug^R^). Treatment with the primary drug in this case activates a sparse network that does not trigger coordinated cell state transitions, but instead drives the surviving cells into multiple distinct drug-resistant states, which may be sensitive to secondary interventions (e.g., siRNA).

Sequential treatments with pitavastatin followed by vinflunine had synergistic effect on killing of the two PD-GSCs (Figure 7D). Specifically, sequential treatment of pitavastatin followed by vinflunine resulted in 5.92 and 1.6 fold-decrease of IC_50_, compared to vinflunine treatment alone (Figure 7D), in SN520 and SN503, respectively. The relative efficacy of sequential treatment with the two-drug combination varied significantly across other PD-GSCs (Supplementary Table S14), with the combination being more effective on pitavastatin responder (SN533 and SN575) than non-responder cells (SN517 and SN521) (Supplementary Figure S14). The poor efficacy of vinflunine on SN503 and other non-responder PD-GSCs is likely because pitavastatin did not induce a coordinated response that places the cells in a vulnerable state from which we predicted the utility of vinflunine based on the transcriptional network. Thus, the coordinated cell-state changes induced by pitavastatin killing of susceptible cells in the responder PD-GSCs pushed the surviving cells along PMT trajectories with generic and patient-specific components, thereby exposing novel vulnerabilities that significantly potentiated net cell killing by sequential treatment with vinflunine.

## DISCUSSION

Inherent plasticity and heterogeneity of GSCs are implicated as underlying reasons for the high rate of GBM recurrence, which often manifest as an even more aggressive and drug-resistant MES subtype (*8–10*). Understanding the mechanisms of primary resistance and trajectories along which GSCs undergo adaptive subtype transitions to acquire resistance are both critical for formulating treatment regimens to prevent recurrence of aggressive and drug resistant GBM (*7*, *68*). In this study, we report five main findings that shed insight into the underlying mechanisms of phenotypic plasticity of PD-GSCs: 1) distinct population structures distinguished two PD-GSCs with acquired (SN520) and primary (SN503) resistance phenotypes, 2) distinct TF network topologies were associated with the two GSC phenotypes, 3) TF network topology was a key determinant of treatment-induced change in the population structure of PD-GSCs, 4) TF network topology inferred from scRNA-seq enabled predictions of underlying mechanistic drivers of primary and acquired resistance, including response trajectories, 5) disruption of primary resistance potentiated killing of non-responder PD-GSCs, and 6) treatment-induced trajectories through which PD-GSCs acquired resistance, exposed vulnerabilities to sequential interventions (siRNA KD of TFs and a secondary drug) targeting transcriptional programs mechanistically associated with cell state transitions.Gene

Primary resistance of SN503 was likely due to a larger pre-existing subpopulation of MES subtype cells, identified by both scRNA-seq and flow cytometry (Figure 2C-F), with elevated expression of OXPHOS and fatty acid metabolism (Figure 5E) and high activity of WNT/β-catenin signaling pathway genes in Pr_503_-18 (Figure 4F) (*7*, *69*, *70*). Hence, pitavastatin treatment was less effective on SN503 and failed to trigger a coordinated transcriptional response across the population of surviving cells in this PD-GSC. By contrast, a smaller proportion of SN520 cells were of the MES subtype (Figure 2C, D) and activity of programs associated with known treatment-resistance mechanisms was low. As a result, pitavastatin killed most SN520 cells, triggering coordinated transcriptional responses across the surviving PD-GSCs, driving their transition into a MES subtype cell state that was > 5-fold resistant to pitavastatin (Figure 2I). Flow cytometry using apoptosis and cell subtype markers; CNV inference; and theoretical calculations based on cell division rates all demonstrated that pitavastatin-induced cell state and phenotypic transitions were mediated by epigenetic mechanisms and not clonal selection. Further, the core TF-TF networks inferred from scSYGNAL analysis were determined by RACIPE simulations as sufficient to generate the observed heterogeneity and treatment-induced cell state changes of the two PD-GSCs. Our findings showed that the TF-TF network topology was likely a key factor in determining the trajectory and potential endpoint(s) of cell-state transitions in response to drug treatment or perturbation. The sparse network of SN503 generated multiple resistant states that were distinct from each other. The interconnected network of SN520, by contrast, generated a gradient of cell states along a PN-to-MES axis offering a plausible explanation as to why GSCs manifest a gradient of resistant states across a range of drugs (*9*). Our findings provide novel perspective on how patient-to-patient variation in the roles of TFs and the topology of their interactions can have profound consequences in driving PMT, likely influencing the rate of GBM progression, recurrence, and metastasis as tumors of MES subtype (*27*, *71*).

By killing a large proportion of cells, pitavastatin treatment triggered a core network of TFs to act sequentially and drive coordinated cell-state transitions across the surviving population of SN520. In so doing, pitavastatin treatment may have generated a bottleneck effect by channeling the surviving SN520 cells along few trajectories, thereby transiently exposing vulnerabilities in associated transcriptional programs across a large segment of those surviving cells, before they transitioned to the MES subtype and acquired a drug-resistant phenotype. Similar constraining effects on GSC plasticity, i.e., fewer cell-state transitions have been observed and attributed to hypoxic micro-environments, unlike the larger number of stochastic cell state transitions that occur under normoxic conditions (*72*). Our findings demonstrate that such constraints on plasticity makes the GSC population less heterogeneous and more vulnerable to siRNAs and drugs targeting transiently activated programs that mechanistically coordinate the cell state transitions. Taken together, these results suggest that the bottleneck effect generated by drug treatment can be exploited to minimize or prevent drug-induced transitions and therapy escape of GSCs.

Notably, the timing of the secondary intervention was critical, with efficacy of potentiation observed only *after* cell-state transitions had been triggered by pitavastatin treatment. The combinatorial interventions (siRNA) were ineffective when each drug-siRNA pair was administered concurrently (data not shown). These findings illustrate the importance of tailoring not just the specific combination of drugs, but also the order and timing of longitudinal treatment schedules based on mechanistic understanding of the causal sequence of events targeted by each individual intervention. Similar benefits from modeling cell state transitions and characterizing trajectories have also been reported in PDGF-driven GBM mouse models. Specifically, the integration of mathematical models that account for the presence of radiosensitive and radioresistant tumor cell states as well as the rate at which state transitions occurred led to an optimized radiotherapy scheduling that improved survival rates of mice (*73*, *74*).

Sequential treatment with vinflunine was effective to varying degrees across other PD-GSCs that were also sensitive to pitavastatin (SN533 and SN575), but was significantly less effective in pitavastatin-resistant PD-GSCs (SN503, SN517 and SN521). This finding suggests that cytotoxic effects were important to expose vulnerabilities, and that the mechanism of killing by pitavastatin and resulting trajectories of escape were likely similar across some of these PD-GSCs. However, variable susceptibilities of PD-GSCs to vinflunine explain why an *N = 1* approach is necessary to uncover patient-specific characteristics and tailor regimen (specific drugs and dosing schedule) to the unique PMT trajectories for each patient (Supplementary Figure S15, Fedele et al., 2019).

The partial generalizability of pitavastatin-vinflunine sequential treatment to other pitavastatin-sensitive PD-GSCs, further suggests that subgroups of patients might share transcriptional regulatory network topologies that drive their tumor cell state transitions along similar trajectories. If this hypothesis is confirmed by analyzing a larger number of PD-GSCs across a diverse range of drug treatments, then stratifying patients based on similar network topologies, instead of steady states of tumor cells, may identify a finite number of topology-matched combinatorial interventions for personalized treatment of most patients (*2*, *3*, *75*).

The causal and mechanistic regulatory influences captured at single-cell resolution in the scSYGNAL network provides a generalizable approach for formulating *N = 1* patient-tailored drug regimens and treatment schedules. Remarkably, we discovered that more than the composition of initial tumor cell states, mechanistic understanding of the topology of the core TF-TF network and its associated dynamics of driving cell state transitions is essential for rationally tailoring sequential treatment regimen to an individual patient. This perspective, borne from these findings, complements prior and current efforts that aim to create frameworks that quantify the hierarchical and multi-state switching that underlie intratumoral heterogeneity in GBM using methods such as Markov chain models or exploratory adaptation models (*76*, *77*). While these approaches define *what* states are present and the probability of transitioning from one state to another, our approach provides mechanistic insights into *how* GSCs are able to navigate the phenotypic landscape (Figure 7E).

Broadly speaking, our findings provide a mechanistic framework for connecting two aspects of phenotypic plasticity of tumor cells, one that characterizes discrete states (*75*), and the second that characterizes cell state continuums, including gradients defined by a neuronal developmental–injury response axis (*78*) or a PN–MES axis (*11*, *79*). Such a framework, like the seminal GBM molecular subtype classification scheme (*2*), will enable integration of the genomic, transcriptomic, and epigenomic landscapes and associated factors that underlie phenotypic plasticity of GSCs and differentiated tumor cells that define intra– and inter-tumoral heterogeneity in GBM (*2*, *4*, *75*, *80*). Ultimately, a systems approach that connects intrinsic regulatory mechanisms wiRAth extrinsic factors, including drug treatment, tumor microenvironment (*72*), and the immune response (*81*), governing phenotypic plasticity of GSCs in an individual patient’s cancer, will be needed for formulating treatment strategies to prevent recurrence of drug-resistant GBM tumors.

## METHODS

### Ethics Statement

Use of human tissue was reviewed and approved by the WIRB-Copernicus Group Institutional Review Board (WCG® IRB). All participants provided written informed consent according to IRB guidelines prior to participation in the study. Only tissue specimens deemed non-essential for diagnostic purposes and that would otherwise be discarded were collected for research purposes.

### Patient samples and patient-derived GBM stem-like cell enrichment

Tumors were obtained from surgeries performed at Swedish Medical Center (Seattle, WA) according to institutional guidelines. Patient samples used in this study were diagnosed as WHO grade IV glioblastoma. GSC cultures were established from freshly resected tumor tissues. Tissue samples were minced into 1mm^3^ fragments and digested with Accutase (Sigma) at 37°C for 15-20 minutes. Neurobasal-A medium (NBM) was added to quench Accutase activity and cell suspensions were filtered through 70μm nylon mesh, centrifuged at 1K rpm for 5 min, resuspended in fresh NBM, and cultured in T75 flasks pre-treated with a laminin solution (1:100 Sigma), which includes incubation of the flasks with the laminin solution at 37°C for a minimum of 30 minutes. PD-GSCs were maintained in NBM with B-27 serum-free supplement, 20 ng/mL EGF, 20 ng/mL FGF-2, 20 ng/mL insulin, 1 mM sodium pyruvate, 2 mM L-glutamine and 1% Antibiotic-Antimycotic.

### PD-GSC in vitro cultures

PD-GSC adherent monolayer cultures were used for all pitavastatin and pitavastatin/vinflunine treatments. Monolayer cultures were maintained in T75 flasks (cell expansion), T25 flasks (pitavastatin-treatment), or 96 well plates (IC_50_ studies) pre-treated with a laminin solution (1:100; Sigma) and incubated at 37°C for a minimum of 30 min. Serum-free culture media consisted of Neurobasal Medium-A (Gibco^TM^) with 2.0% (v/v) B-27 serum-free supplement minus vitamin A (Gibco^TM^), 20 ng/mL EGF (PeproTech Inc.), 20 ng/mL FGF-2 (PeproTech Inc.), 20 ng/mL insulin (Sigma), 1 mM sodium pyruvate (Corning), 2 mM L-glutamine (Gibco^TM^) and 1% Antibiotic-Antimycotic (Gibco^TM^). PD-GSC monolayer cultures were maintained at 37°C, 5% CO2 atmospheric oxygen with culture pH monitored with the phenol red. Cultures were refed every 2-3 days. PD-GSC cultures tested were within 10 passages from the initial GSC enrichment from the original tumor biopsy.

PD-GSCs were passaged by dissociating monolayer cultures from the respective substrate by treating the cells with the dissociation reagent Accutase (1mL/25cm^2^) or TrypLE^TM^ (1mL/25cm^2^ – see *Flow cytometry CD44 and CD133 analysis* section) at 37°C for 5min. Pre-warmed (37°C) serum-free culture media (described above) was then added to quench dissociation reagent activity (1:3 media:dissociation reagent ratio). The resulting cell suspension was centrifuged at 1K rpm (193g) for five minutes. The cell pellet was resuspended in fresh serum-free culture media, and added to QS serum-free culture media in a new laminin-treated flask. Final culture volumes were as follows: T75 – 10mL, T25 – 5mL, 96-well plate – 100μL. Laminin treatment involved incubating flasks (or 96 well plates) with a laminin working solution (5mL/75cm^2^), which consisted of stock laminin (Sigma) diluted 1:100 in phosphate buffer solution, at 37°C for a minimum of 30 min.

### Flow cytometry – apoptosis, caspase 3/7-mediated apoptosis, and cell-death

Data acquisition of surface protein markers was performed on the Attune NxT Flow Cytometer (ThermoFisher Scientific). PD-GSCs were dissociated from their respective substrate using Accutase and washed twice with PBS + FBS serum (10%), which involved centrifugation at 1K rpm (193g) for 5 min, supernatant removal, and cell pellet resuspension with the PBS + FBS serum (10%). The supernatant wash was removed and the cell pellet resuspended in the PBS/FBS solution to the desired concentration of 1e6 cells/mL. To assess apoptosis, caspase 3/7-mediated apoptosis, and cell death within the GSC populations, cells were stained with Annexin V conjugated with Alexa Fluro 568 (Invitrogen A13202), CellEvent^TM^ Caspase 3/7 detection reagent (Invitrogen C10423), and SYTOX^TM^ AAdvanced Dead Cell Stain (Invitrogen S10349), simultaneously. Samples were stained following each of the manufacturer’s protocol, respectively. Gating for positive and negative expressing cells was performed using FlowJo V10 based on multiple controls including, 1) unstained negative controls, 2) heat-inactivated cells (incubated in a 60°C water bath for 15 min), which served as positive controls for apoptotic and dead cells, and 3) Fluorescence minus one (FMO) controls to define an upper boundary for background signal on the omitted signal and gate for positively stained populations in multi-color experiments.

### Flow cytometry – CD44 and CD133 analysis

Samples from each treatment condition were collected using TrypLE^TM^ (Gibco^TM^) to dissociate and remove the cells from the culture flasks. TrypLE^TM^ (1mL/25cm^2^) was used to minimize any structural changes on CD44 and CD133 surface proteins during the dissociation process (*82*). Subsequent sample processing prior to antibody staining was identical to how samples were processed for apoptosis, caspase 3/7-mediated apoptosis, and cell-death cytometry assessment. An anti-Hu CD44 antibody conjugated with PE (eBiosciences^TM^) and an anti-Hu/Mo CD133 antibody conjugated with FITC (eBiosciences^TM^) were used to assess expression of these two surface proteins across each PD-GSC population. Samples were simultaneously treated with both antibodies per vendors’ recommendations. Analysis of flow cytometry data was performed using FlowJo V10. Fluorescent signal gating was set based on multiple control samples including: 1) unstained PD-GSC negative controls, 2) vendor-recommended isotype controls (Mouse IgG1 kappa isotype and Rat IgG2b kappa isotype for anti-Hu CD133 and anti-Hu/Mo CD44, respectively, 3) human GBM stem cells (Cellprogen Inc.), which served as a positive control cell line for both CD133 and CD44 (per vendor’s specification), and 3) Caco2 cells, (ATCC) which served as a positive control cells for CD133 and negative controls for CD44.

### Pitavastatin treatment of PD-GSCs for scRNA-seq and flow cytometry analysis

PD-GSCs were incubated in serum-free culture media (described above) with pitavastatin (6μM). Stock pitvastatin calcium (Selleck Chemicals LLC) was dissolved in DMSO to obtain a stock concentration of 10mg/mL and stored in aliquots at –80°C. Stock pitavastatin calcium solution was serially diluted in serum-free culture media to 100μM and then to the final concentration of 6μM with a final DMSO concentration of 0.053% (v/v).

To monitor longitudinally PD-GSC response to pitavastatin, we performed a reverse time-course treatment by adding pitavastatin to SN520 and SN503 cultures in a staggered fashion such that the longest (4-day) treatment would have drug added first. Subsequent addition of pitavastatin would occur on following days for 3– and 2-day treatment, respectively. This reverse time course design allowed us to collect all samples simultaneously on day four following the initial addition of pitavastatin. Because pitavastatin was added to PD-GSCs on different days, flasks were inoculated at slightly different cell densities to account for cell growth that would occur in between inoculation and time of pitavastatin addition. Consequently, scRNA-seq library preparation of all samples for a particular PD-GSC population occurred simultaneously to minimize batch effects due to individual sample processing (Supplementary Table S15)

Prior to T25 flask (BioLite^TM^) inoculation for pitavastatin treatment, PD-GSCs were first expanded in a T75 flask (BioLite^TM^). Once the culture was confluent, the culture was harvested and split into laminin-treated T25 flasks. Upon inoculation, cells were incubated in serum-free culture media at 37°C for 24 hours to allow cells to adhere to the interior surface of the flask. Following the first 24 hours, serum-free culture media was replaced with serum-free culture media with pitavastatin (6μM) in T25 flasks predetermined to receive a 4-day treatment. Spent culture media would then be replaced with fresh culture media with pitavastatin (6μM) on subsequent days for D3 and D2 treatment conditions.

Upon the completion of the 4-day treatment, spent media was removed and cells were harvested using Accutase^TM^ (1mL/25cm^2^). To prevent any cell-free DNA/RNA from treatment-induced lysed cells contaminating single-cell samples, we first processed a portion of the cell harvest solution using the dead cell removal kit (Miltenyi Biotec 130-090-101) to remove any cell debris to avoid any free RNA from lysed cells from getting mixed in with mRNA to be extracted from live cells. Samples were processed per vendor’s specifications. The result was a cell suspension of the remaining live cells post vehicle– or pitavastatin-treatment. Cell suspension was then processed for scRNA-seq profiling per the 10X Chromium platform.

### scRNA-seq library prep and sequencing

Single-cell RNA sequencing was performed using the 10X Chromium v2 system. Library preparation was performed using 10x manufacturer instructions on an Illumina NovaSeq 6000. scATAC-seq was performed as per manufacturer instructions (Single-cell ATAC Reagent Kits v1.1 UserGuide RevD) and sequenced on an Illumina NextSeq 500.

### Multi-passage, pitavastatin treatment

PD-GSCs were harvested from a T75 flask and passaged into replicate T75 flasks for either pitavastatin (6μM) or vehicle (DMSO) treatment (2e6 cells/flask). Concomitantly, a portion of those PD-GSCs were used to inoculate laminin-treated 96 well plates for drug-dosing analysis (see *IC*_50_ *Analysis* section). On D4, PD-GSCs were harvested using Accutase (1mL/25cm^2^) as described previously. Cell suspensions were spun at 1000rpm (193g) for five minutes. Cell pellets were then resuspended with serum-free culture media (200,000 cells/mL) to inoculate 96 well plates (100μL/well, 20,000 cells/well) for subsequent IC_50_ determination. PD-GSCs were incubated in serum-free culture media in 96 well plates for 48 hours to allow for cell attachment prior to replacing spent media with serum-free media with pitavastatin (or vehicle). Treated cells were incubated at 37°C for four days. Following the four-day treatment, cell viability was measured via MTT assay as described below.

### DNA quantification via propidium iodide (PI) staining

PD-GSC cultures were treated with pitavastatin (or vehicle control) in a reverse time-course manner as described previously (*Pitavastatin treatment of PD-GSCs for scRNA-seq and flow cytometry analysis* section). Following cell harvest, PD-GSCs were washed with PBS and spun down at 1000 RPMs (193 g) for 5 minutes. PD-GSCs were then fixed with cold 70% ethanol by adding 70% ethanol drop-wise to the pellet while vortexing. Cells were fixed in 70% ethanol overnight at 4°C. Once fixation was complete, the PD-GSCs were washed twice in PBS, spun down at 1000 rpms for five minutes with careful removal of the supernatant so as to avoid any cell loss. PD-GSCs were then treated with 50μL of ribonuclease (100μg/mL stock) to remove any RNA and ensure only DNA would be stained. Finally, 200μL of propidium iodide (PI, 50μg/mL stock) was added to the fixed and treated cells prior to flow cytometry analysis.

### IC_50_ Analysis and MTT viability assay

3-(4,5-Dimethyl-2-thiazolyl)-2,5-diphenyl-2H-tetrazolium bromide, (MTT) assay was used to determine the effects of pitavastatin on the viability of the non-responsive and responsive GSC populations. Briefly, 20,000 cells/well were plated in laminin-treated 96-well plates with 100uL of culture media. Following an initial 24hr incubation, the cells were treated with 100μL of culture media with pitavastatin at varying concentrations (0.0, 0.1, 0.6, 1.0, 3.0, 6.0, 10.0, 33.0μM) and incubated at 37C for four days. Vehicle amounts were adjusted such that the vehicle concentration in all conditions was equivalent to the maximum drug dosage tested (DMSO 0.2% v/v). Following the 4-day treatment, spent media was replaced with 100μL of serum-free culture media with MTT (0.5mg/mL) and incubated at 37°C for 60 minutes. Following incubation, supernatant from each well was discarded and replaced with 100μL of DMSO to dissolve the formazan crystals formed during MTT incubation. Absorbance (*A_i_*, where *i* is the drug concentration) was measured via spectrophotometer at 570nm (Synergy H4, Agilent Technologies, Inc.). Relative viability was calculated using the following formula: relative viability = *(A_i_ – A_background_)/A_0.0_ * 100%*, where *A_background_* is the absorbance from DMSO. IC_50_ values were calculated by using a 4-parameter log-logistic model determined by the *drm()* function within the *drc* package in R. Here, the upper limit of the log-logistic model was set to 100%.

### siRNA treatment

Following a 24hr incubation period, cells were treated with 5μM of Accell SMARTpool siRNA or Accell SMARTpool Non-Targeting siRNA (Dharmacon Inc.). Lyophilized SMARTpool siRNAs were resuspended in 1X siRNA buffer (Dharmacon Inc.) and subsequently diluted in serum-free culture media to a final concentration of 5μM. Based on vendor recommendations, Accell siRNA designs facilitate siRNA delivery to the target cell and do not require additional transfection reagents. Accell SMARTpool siRNAs pools consist of four separate siRNAs designed to target a particular gene. To test the efficacy of siRNA-targeted knockdown of specific TFs, siRNA (5uM) and pitavastatin (1.0μM or 6.0μM for SN520 and SN503, respectively) were added simultaneously followed by a four-day incubation at 37°C.

### Bulk RNA-seq library prep and sequencing

Total RNA was extracted from PD-GSC cultures using mirVANA^TM^ miRNA isolation kit (ThermoFisher). Residual DNA was removed using the RQI RNAse-Free DNase kit (Promega). Total RNA was then quantified using the Agilent RNA 6000 nano kit (catalogue number) on the Agilent 2100 BioAnalyzer. 1μg of of high purity RNA was used as input to the Illumina TrueSeq Stranded mRNA Library Prep Kit and sample libraries were generated per manufacturer’s specifications. The RNA-seq libraries were sequenced on the NextSeq 500 next gen sequencer using a paired end high-output 150bp v2.5 flowcell. Sequence intensity files were generated on instrument using the Illumina Real Time Analysis software. The resulting intensity files were de-multiplexed with the bcl2fastq2 software.

### Processing and normalization of bulk RNA-seq data

Raw RNA-seq data of samples encoded in FASTQ-files were subjected to a standardized RNAseq alignment pipeline. In summary, RNA-seq reads were trimmed and clipped of Illumina sequence adapters via Trim Galore (https://github.com/FelixKrueger/TrimGalore), mapped to human reference genome (GRCh38) using STAR (v2.7.3a), and counted using HTSeq (v 0.11.1). Individual sample counts were combined into a single data object using the *DESeqDataSetFromHTSeqCoun*t function in DESeq2 (*83*). Sample-specific size factors were determined and used to normalize counts, which were transformed using regularized log transformation for subsequent downstream analysis, performed in R.

### scRNA-seq data QC filtering and normalization

We initially processed the 10X Genomics raw data using Cell Ranger Single-Cell Software Suite (release 3.1.0) to perform alignment, filtering, barcode counting, and UMI counting. Reads were aligned to the GRCh38 reference genome using the pre-built annotation package download from the 10X Genomics website. We then aggregated the outputs from different lanes using the *cellrange aggr* function with default parameter settings.

SN520 and SN503 scRNA-seq data sets were QC-filtered separately prior subsequent downstream analysis. To minimize inclusion of poor-quality genes and single-cell samples per sample set, we applied the following QC filters: 1) mitochondrial genes must comprise ≤ 6.5% of the number of uniquely mapped genes/cell, and 2) total counts/cell should be ≥ 7500 and ≤ 60,000. Post QC-filtering, each scRNA-seq data set included: 5,402 cells expressing up to 18,227 genes (SN520) and 5,722 cells expressing up to 18,797 genes (SN503). Subsequent normalization and downstream analysis (e.g., DEG and functional enrichment analysis) was performed using the Seurat v3.2.2 platform (*84*).

Normalization was performed for each scRNA-seq dataset separately by computing pool-based size factors that were subsequently deconvolved to obtain cell-based size factors using the *computeSumFactors* function within the *scran* package (version 1.10.2) (*85*) in R. Normalized log expression values were used for subsequent downstream analysis.

### Batch integration of scRNA-seq data

As each PD-GSC-specific data set was collected separately, we performed batch correction on the scRNA-seq data to integrate the SN520 and SN503 data sets by applying the Harmony algorithm (*32*). Subsequent SNN-graph formation and UMAP embedding was performed on the Harmony-corrected PCs (Fig. 1E).

### Cell-cycle analysis

To annotate individual cells with their respective cell cycle phase, we performed cell cycle analysis using the Seurat program. Briefly, core sets of 43 and 54 genes associated with the S– and G2/M– phases, included in the Seurat platform, were used to determine a cell-cycle phase score based on the expression of the respective markers. Based on these scores, cells were assigned to be either in G1 or G2/M phase. Cells not expressing genes from either set were considered as not cycling and assigned to the G1 phase. Using these quantitative scores, we also regressed out cell-cycle effects on expression for each cell using the *ScaleData* function in Seurat as part of the pre-processing steps to QC the scRNA-seq data.

### Cluster identification and analysis of differentially expressed genes (DEGs)

After quality control and filtering the scran-normalized scRNA-seq data, we performed dimensionality reduction via principal component analysis (PCA). The first 30 principal components were used as a basis to create a shared nearest neighbor (SNN) graph of the single-cell samples. From this graph, clusters of single cells were identified via Louvain clustering of nodes, i.e., single cells, from the SNN graph.

To identify DEGs in each of the SNN-clusters identified across the primary tumor and PDX single-cell samples, the *FindMarkers* function in Seurat was used. In particular, the Wilcoxon rank sum test was used with the following cutoff values to identify DEGs: absolute log-fold change ≥ log2(1.5), with a minimum proportion of 10% of the cells of interest expressing the gene of interest, and an FDR-adjusted p-value ≤ 0.1.

### Gene set variance analysis (GSVA) enrichment scores and statistical significance

Gene set variance analysis GSVA (version 1.34.0, R package) (*31*) was used to determine enrichment scores of GBM molecular subtypes. To define the dominant molecular subtype gene expression signature in each single cell, we used an amalgamation of the original gene sets that defined the classical, proneural, and mesenchymal subtypes (*2*) and refined molecular subtype gene sets (*3*) for GSVA.

### Critical Transition Index (***I_c_***)

A brief explanation of *I_c_* from (*52*) is reproduced for reference. The critical transition index is a scalar value that quantifies if a cell is undergoing (high Ic) or has undergone some critical transition and reached some stable cell state (low Ic). *I_c_* is calculated according to the following:

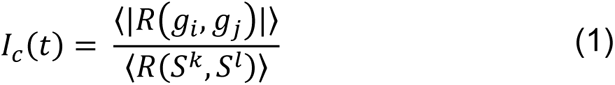

Where *R* is Pearson’s correlation coefficient between two observed cell state vectors *S^k^* and *S^l^* or between two “gene” vectors *g_i_* and *g_j_*, respectively, taken from the gene expression data matrix representing the state(s) of a “cell ensemble” *X(t)*

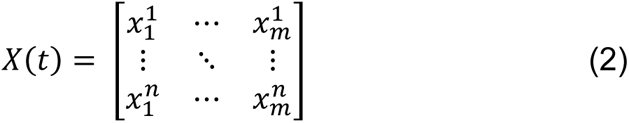

*X(t)* thus represents the data of a “measurement point”, with access to finer-grained layer of information given the single-cell nature of the data. Each row represents a single-cell in some state *k* within the cell-ensemble of *n*-cells in *m*-dimensional gene state space – 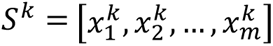. Each column represents gene *i’s* expression across *n* cells from said “cell ensemble” *X(t)*, where 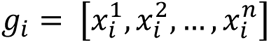. The brackets 〈⋯ 〉 in equation 1 represent the average of all correlation coefficients *R* between all pairs of state vectors *S* or gene vectors *g* from matrix *X(t)*. Here, a cell-ensemble represented the population of PD-GSCs at a particular treatment time-point (D0, D2, D3, or D4).

The underlying premise is that cells that have undergone some critical transition into an attractor state will be nominally expressing the same distinct gene expression pattern, with the exception of deviations due to stochastic fluctuations. Consequently, cells of the same differentiated state will be expressing similar gene expression programs and will correlate highly with one another. Characteristic gene expression of cells within a particular attractor state is affected by symmetric random fluctuations. Thus, gene-to-gene coupling is dominated by noise, reducing gene-to-gene correlations. Conversely, destabilized cells undergoing some transition, requires some non-random shift in gene expression patterns that override the symmetric noise expected in cells within a stable attractor state.

### MINER network inference

An additional gene-filtering step was performed on the QC scRNA-seq data sets to identify a common gene set between SN520 and SN503 – only common genes having a minimum gene count ≥ 2 in a minimum of 20 cells were considered for network inference. This resulted in a common gene set of 9,089 common genes used in SN520 and SN503 for MINER3 network inference.

To infer regulons within single cells, we applied the MINER (*86*) workflow to the SN520 and SN503 scRNA-seq data sets independently. As part of the scSYGNAL framework, the MINER algorithm involves a suite of functions that enables the inference of causal mechanistic relationships linking genetic mutations to transcriptional regulation. Because our datasets did not include any mutational profiling, we primarily focused on identifying regulons, based on co-expression clustering and enrichment of transcription factor binding motifs present in those co-expression clusters identified, and calculated the activity of these regulons in the single-cell samples. Broadly speaking, regulon activity represents the “eigengene” value in an individual cell. Regulons are identified, in part, by performing PCA on the normalized scRNA-seq data profiles to identify principal components in which decreasing amounts of variation across genes are captured along each principal component – defined as a linear combination of gene expression values. This linear combination of weighted gene expression values defines the eigengene value per sample (*41*, *42*, *86*, *87*). Alternatively, the eigengene is defined as the first principal component of the module expression matrix composed of expression values of regulon genes across samples. It is a scalar representation of expression of gene members for a regulon in an individual sample (*87*).

To determine the significance of each inferred regulon, we performed a permutation test to determine the possibility of obtaining an eigenvalue corresponding to the first principal component of a regulon (across all single-cells) of equal or greater value. The eigenvalue represents a summarizing value of all the genes in the regulon, i.e., eigengene and thus if these genes are indeed coregulated or are correlated, the eigengene value would be higher than that of randomly selected set of genes. Next, we randomly select a set of genes having the same number of members as the original regulon and calculate the corresponding eigengene value for the permuted regulon. This procedure was repeated 1,000 times to create a null distribution of eigengene values. We repeated this procedure for each inferred regulon. Those regulons whose eigengene values were greater than the 95th percentile of their respective null distribution were considered significant. These eigengene values represented regulon “activity” within each cell. We further filtered out regulons in which the first principal component from the module expression matrix composed of expression values of regulon genes across samples did not account for at least 20% of the variation of the module expression matrix. From these two criteria, statistical significance of an eigengene and variance explained within the module expression matrix were used to refine the number of regulons to include for SN520 and SN503, respectively.

### Pseudotime analysis

We applied Monocle v3 in R (*88*, *89*) to organize cells along a pseudotime axis and identify distinct trajectories along which transcriptomic expression states putatively transition. Scran-normalized scRNA-seq datasets were used to infer pseudotime trajectories for SN520 and SN503 independently using the *learn_graph* and *order_cells* function in Monocle v3 (v1.2.7) and default parameter settings.

### Locally estimated scatterplot smoothing (LOESS) regression analysis

We performed LOESS regression on individual TF expression across the single cells along the inferred pseudotime trajectories. This allowed us to fit a polynomial regression line through the highly variable single-cell gene expression to identify any underlying patterns that may be present over pseudotime. LOESS regression of normalized single-cell gene expression along pseudotime was performed using the *loess* function within the *stats v3.6.2* package in R.

### TF-TF network topology inference

To generate TF-TF network topologies, we cross-referenced all regulator-target gene connections inferred by MINER3 against the transcription factor binding site database (tfbsdb.systemsbiology.net), focusing on only those interactions that involved pairs of TFs that were also regulators for some regulon. The type of TF-TF interaction was determined by the sign of the pairwise Pearson correlation between the two components – positive correlations were interpreted as activating interactions while negative correlations were interpreted as inhibiting interactions. We further refined the TF-TF network by removing those interactions having an absolute Pearson correlation coefficient *(r)* below a statistically significant minimum threshold, determined by permutation analysis (|*r* | ≥ 0.17 for SN520 and |*r* | ≥ 0.16 from SN503). Permutation tests involved randomly mixing expression values across genes within a single-cell and calculating Pearson’s *r* among all gene pairs across all PD-GSCs for SN520 and SN503 independently. This process was repeated 1000 times to create a null distribution of Pearson correlation coefficients.

To determine the statistical significance of each network TF-TF network topology, we performed two sets of permutation tests (Supplementary). Briefly, the first set of permutation testes involved permuting the network topology, where node labels and edges were permuted such that the number of edges and nodes remained consistent, we performed dynamic simulation for the permuted network using initial condition, i.e., TF expression profiles from a randomly selected untreated (D0) cell for each PD-GSC, respectively. The simulated results were then compared to experimental data to determine cosine similarity values. This permutation-simulation-comparison process was repeated 1,000 times to create a null distribution of cosine similarity values. The distribution of cosine similarity values derived from the original TF-TF network topologies were significantly higher than the permuted similarity values (SN503 empirical p-value = XXX, SN520 empirical p-value = YYY). The second set of permutations involved permuting the gene expression data, mixing the gene and cell ids to see if similar TF-expression states could be achieved by random chance. Cell and gene labels were permuted 1000 times to create a permuted distribution of TF-expression states, which were then compared to the original experimental states, defined by hierarchical clustering, using pairwise cosine similarity values (Supplementary Figure S10).

### RACIPE simulations

Simulations were performed using the sRACIPE package v1.16.0 in R. Briefly, using sRACIPE we generated an ensemble of ordinary differential equation (ODE) models based on associated chemical rate equations with distinct, randomly generated kinetic parameter sets. From the ensemble of models, we analyze the resulting distribution of steady states and identify robust phenotypes supported by the core TF network. The inferred TF-TF network topology for SN520 (or SN503) was used as the input circuit for the *sracipeSimulate* function. An integral step size of 0.2 and simulation time of 100 was used for simulations.

To verify the ability of the network topology to recapitulate observed TF expression states, we initialized the network by randomly selecting 1,000 expression profiles (with replacement) for the respective TFs from D0 scRNA-seq profiles for each PD-GSC, i.e., initial conditions that were paired with 1,000 parameter models randomly selected by the *sracipeSimulate* function (default settings used).

To explore the plausible network states supported by each network topology, we initialized each network topology by using 100 randomly selected initial conditions that were used across 10,000 randomly selected parameter sets, which resulted in an ensemble of 1 million simulated steady-states. To determine the dominant steady states from the ensemble of simulations, all Euclidean pairwise distances were calculated. Those simulated states that had a Euclidean pairwise distance ≥ 4.0 (scSYGNAL-520) or ≥ 1.92 (scSYGNAL-503) were labeled as a “non-redundant” state. The distance thresholds were found to be the ≥ 99^th^ percentile of permuted Euclidean pairwise distances for each PD-GSC, which was determined by randomly selecting 1,000 pairs of simulated states and calculating all pairwise Euclidean distances. This process was repeated 10 times to create a distribution of 10 million pairwise Euclidean distances. From these distance thresholds, we identified 6,519 (scSYGNAL-520) and 4,223 (scSYGNAL-503) simulated states were deemed as unique states. We then hierarchically clustered each set of distinct, “non-redundant” states and identified four dominant states that were supported by each TF-TF network topology (Figure 6C, E). To classify a “redundant” simulated state, we assigned it the same state as its nearest “non-redundant” neighbor, based on Euclidean distance.

### RACIPE convergence tests

To verify that the number of initial conditions and parameter sets would sufficiently converge to steady state solutions across the initial condition and parameter space, we performed a series of simulations using 100 randomly selected initial conditions across different number of model parameters (1e3, 2e3, 4e3, 6e3, 8e3, and 1e4). The result was a series of simulations consisting of six different ensembles of simulated states, one for each model parameter set, with each ensemble associated with a randomly selected set of initial conditions. This series of simulations was performed in triplicate. For each set of results, we identified the unique states using the same Euclidean distance thresholds described in *RACIPE simulations*. Next, we determined the Kullback-Liebler (KL) divergence for these simulated states across the triplicate set of simulations for each set of results (Supplementary Figure S11).

### Random Forest analysis of RACIPE simulations

Random forest analysis was performed on RACIPE simulations, i.e., simulated transcriptional states for SN520 and SN503 using *randomForest* function (default parameters) from the *randomForest* package v4.7-1.1. Simulated state classifiers were based on hierarchical clustering of the unique (non-redundant) simulated states as described in *RACIPE simulations*.

### Drug Matching Identification

To identify drugs targeting elements within the transcriptional programs identified from the network analysis, we applied the Open Targets platform tool (https://www.targetvalidation.org/). The platform integrates a variety of data and evidence from genetics, genomics, transcriptomics, drug, animal models, and literature to score and rank target-disease associations for drug target identification. We focused our search on identifying drug-target matches for only those drugs associated with any cancer treatments that had reached Phase IV matching with regulon genes associated with SN520. Originally, 28 drugs paired with genes across 17 regulons. We further refined the list of potential drug candidates to those drugs associated with GBM, reducing the number of candidate drugs to eight, including vinflunine.

## DATA AND CODE AVAILABILITY

All single-cell RNA-seq data will be deposited in dbGaP. All code is available upon request. Any additional information required to reanalyze the data reported in this paper is available upon request.

## Supporting information

Supplemental material revised

Supplemental tables revised

## ACKNOWLEDGEMENTS

We thank M. Strasser for insightful discussions and advice on simulating and analyzing TF-TF network dynamics; M. Arietta-Ortiz for advice on TF-TF network analysis and testing; H. Hampton on advice on experimental design and analysis of flow cytometry data; A. Akade for helpful discussion on stem-cell culture methodologies; C. Lausted for advice on experimental design for single-cell RNA-seq; and the entire Baliga laboratory for general support and advice. We would also like to thank the ISB Molecular Core for their services in preparing and sequencing the single-cell samples and Timothy J. Martins and the University of Washington Quellos High-throughput Screening Core for advice and services in running HTP screens. J.P. was funded by a fellowship from the NIH (F32-CA247445) and currently supported by NIH grant R01-CA259469. P.H. was funded by the Ben and Catherin Ivy Foundation and is currently supported by R01-CA259469 and philanthropic funding from Swedish Medical Center Foundation. A.L. was supported by R01-AI141953. M.P. is supported by R01-AI128215 and R01-CA259469, R.C. was supported by R01-CA259469, W.W. is supported by R01-AI128215 and R01-CA259469, and S.T. is supported by R01-AI128215 and R01-CA259469. H.L. is supported by R01-CA259469 and philanthropic funding from the Swedish Medical Center Foundation. A.P.P. is supported by R01-NS119650, the Burroughs Wellcome Career Award for Medical Scientists, and Discovery Grant from the Kuni Foundation. C.C. is supported by R01-CA259469. S.H. is supported by R01-GM109964, R01-CA226258, R01-GM135396, and R01-CA255536. N.S.B. is supported by R01-AI128215, R01-CA259469, and R01-AI141953.

## AUTHOR CONTRIBUTIONS

J.P., S.H., and N.S.B. conceived the study. J.P. designed all experiments with guidance from A.L., M.P., P.H., and C.C. Pitavastatin-treatment experiments for bulk and single-cell RNA-seq, and all flow cytometry-related experiments were performed by J.P. M.P. prepared samples for bulk RNA sequencing. J.P., M.P., and R.C. performed all siRNA-related experiments. P.H. and C.C. organized and executed the HTP drug screen. P.H. and H.L. performed sequential drug treatment experiments. J.P. analyzed all data and performed all network dynamics simulations with guidance from A.P.P., S.H., and N.S.B. W.W. and S.T. performed miRNA regulation and drug targeting analysis. J.P. and N.S.B. wrote the paper with input from all authors.

## DECLARATION OF INTERESTS

NSB is a co-founder and member of the Board of Directors of Sygnomics, Inc., which will commercialize the SYGNAL technology. The terms of this arrangement have been reviewed and approved by ISB in accordance with its conflict of interest policy. APP is a consultant for and has an equity interest in Sygnomics, Inc. CC and PH hold a patent titled “Methods and panels of compounds for characterization of glioblastoma multiforme tumors and cancer stem cells thereof” (U.S. Patent No. US11499972B2). JP, ST, SH, and NSB have applied for a patent titled “Cancer drug sensitivity, treatment, and progression determination” (U.S. Patent application number 63/549,391).

## Notes

### Summary of Updates

Figure 1 revised; Figure 6 G-H revised; Figure 7D revised; Declaration of interest updated; Supplemental files and tables updated.

