## Supplemental material revised for "Gene regulatory network topology governs resistance and treatment escape in glioma stem-like cells"

**Title:**

**Affiliations:**

#### Supplementary Text

##### *Selection and proliferation of mesenchymal GSCs does not account for changes in PD-GSC population structure*

Given the mounting evidence supporting the ability of GSCs to undergo cell state transitions in response to drug treatment, exemplified in PMT, it was likely that the shift in the proportion of molecular subtypes observed in SN520 was a result of such a transition. However, it was possible that the shift in molecular subtype was the result of a selection of a pre-existing subpopulation of MES GSCs. To confirm whether changes in population structure occurring in PD-GSCs were due to non-genetic changes in cell state or drug-induced selection, we performed both theoretical calculations and DNA quantification to determine the feasibility of a selection process driving the observed changes in SN520 population structure. Using the population structure and number of cells collected at each time point, we considered three scenarios for our theoretical calculations: 1) only pre-existing MES cells survived treatment with negligible proliferation, 2) only pre-existing MES cells survive and proliferate, and 3) assuming selection was the driving force, what cell doubling time ($t_d$) would be required to produce cell counts observed.

We analyzed multiple scenarios under which a change in population structure could occur: *i)* Selection of MES PD-GSCs, *ii)* selection and proliferation of MES PD-GSCs only, and *iii)* concurrent proliferation of MES PD-GSCs and death of all non-MES PD-GSCs. In all scenarios, cell counts and estimated doubling times ($t_d$) for SN520 were used (Supplementary Figure S6).

In scenario *i)*, the simplest case – we assumed that the initial amount of MES PD-GSCs at D0 (24,657 cells) remained consistent through the 4-day experiment. As we estimated a total of 537,500 total viable cells by the end of the 4-day pitavastatin treatment, the proportion of MES cells would only account for 4.6% of the total PD-GSC population, which differs tremendously from the 94% proportion present in the surviving cells (505, 421 MES PD-GSCs). Alternatively, had a majority of the non-MES PD-GSCs died during treatment, it is theoretically possible that the MES subtype could make up 94% of the surviving cells, if not more. The theoretical final cell counts, however, would not match with experimental results.

In scenario *ii)* we analyzed an extension of scenario *i)* alternative in that it was assumed that all non-MES PD-GSCs were eliminated by the end of the 4-day treatment and that only MES PD-

GSCs grew during this time. To estimate final cell amounts by day 4, we assumed exponential growth, characterized by the following:

$$x_f = x_i * 2^n \quad \text{Eqn. 1}$$

Where $x_f$ is final number of PD-GSCs, $x_i$is the initial number of PD-GSCs, and $n$ is the number of doublings that occurred. Here the number of doublings is equivalent to

$$n = \frac{\Delta t}{t_d} \quad \text{Eqn. 2}$$

Where $\Delta t$ is the duration of the experiment and $t_d$ is the doubling time, which was determined a prior (Supplementary Figure S6). In this case, SN520 had a doubling time of ~90 hrs. Based on Eqns. 1 and 2, the total number of MES cells totaled 51,421 cells, which also falls short of the 537,500 cells counted at day 4.

In scenario *iii)* we determined what $t_d$ would be required of the MES cells to match experimental observations. Based on the initial and final number of MES PD-GSCs, 24,658 and 51,421 cells, respectively, and a 3.94 day duration time, we found that td of 21.9 hr would be required for the initial number of MES PD-GSCs to match what was experimentally measured at day 4. This represents an approximate 4-fold decrease in doubling time, which is highly unlikely. It is important to note that these theoretical calculations represent the ideal or maximal growth rates for the MES PD-GSCs. In other words, despite calculations, which favored MES growth, some other factor or process most likely contributed to the increase in proportions of MES PD-GSCs. These results, taken together with the low number of PD-GSCs in G2/M phase, based on cell cycle annotation and DNA quantification, strongly point towards pitavastatin treatment induced a MES transition in SN520.

***Differential Expression Gene and Clustering Enrichment Analysis***

DEG and enrichment analysis revealed several insights into the cellular response and sequence of responses for each PD-GSC were revealed from DEG analysis. As SN520 expressed a clear coordinated response during treatment, we provide additional details on the DEG and enrichment analyses and results.

SN520 Clustering & Enrichment. Vehicle control cells of SN520 from all time points were evenly distributed across eight Leiden clusters ($cl_{520}$-0, $cl_{520}$-1, $cl_{520}$-3, $cl_{520}$-5, $cl_{520}$-8, $cl_{520}$-10, and $cl_{520}$-11), a majority of which were enriched for genes of oxidative phosphorylation (OXPHOS) and growth-related hallmark pathways like E2F targets and G2M checkpoint (Figure 3E, Supplementary Table S2). Together these findings suggested that, in the absence of drug treatment, SN520 cells proliferated using OXPHOS as a mode of energy production [1,2]. By contrast, five clusters ($cl_{520}$-12, $cl_{520}$-4, $cl_{520}$-7, $cl_{520}$-13, and $cl_{520}$-6) were predominantly enriched with cells from a single time point of pitavastatin-treatment. In fact, Leiden clusters could be organized longitudinally based on the relative proportions of drug-treated cells from each day to recreate the likely sequence of events triggered by pitavastatin (Park et al., 2022) (Figure 3C). For instance, temporal ordering of D2 and D3 pitavastatin-treated cell clusters ($cl_{520}$-2 → $cl_{520}$-9 → $cl_{520}$-4 → $cl_{520}$-7) revealed sequential differential regulation of cholesterol homeostasis, fatty acid metabolism, MTORC1 signaling, a regulator of lipid formation (Tian, Li, & Zhang, 2019), and cholesterol biosynthesis and maintenance. This sequential differential regulation of functions across D2 and D3 cells was consistent with the mechanism of action of pitavastatin (i.e., inhibition of cholesterol biosynthesis). In addition, enrichment of apoptosis ($cl_{520}$-4, $cl_{520}$-6, and $cl_{520}$-7) and TNFα signaling via NFκB ($cl_{520}$-6 and $cl_{520}$-7), with progressively higher proportions of D4 pitavastatin-treated cells suggested a mechanism of killing by pitavastatin. Specifically, the findings showed that on D4, pitavastatin treatment had induced TNFα signaling, which activated apoptosis within a subpopulation of SN520 GSCs and was consistent with both annexin V cytometry results (Supplementary Figure S2) and timing of maximal cell death rate (Figure 1B). Further, the high proportion of D4 pitavastatin-treated cells in $cl_{520}$-6 and $cl_{520}$-7 indicated that the cytotoxic effects resulted in the upregulation of cellular stress responses including unfolded protein response, protein secretion, and p53 pathway.

As described in the main text, $cl_{520}$-6, $cl_{520}$-7, and $cl_{520}$-13, which were all D4 pitavastatin-treated cell clusters, were enriched for both apoptosis and EMT, which aligned with the timing of cell state transition to the MES subtype (Figure 2C, D). These findings were consistent with previous studies that reported that TGFβ can simultaneously induce apoptosis and EMT during tumor formation and progression. Cell fate correlated with cell-cycle phase, with tumor cells in G2/M phase undergoing apoptosis and those in G1/S undergoing EMT [3,4]. As cells in $cl_{520}$-6, $cl_{520}$-7, and $cl_{520}$-13 were in G1/S phase, explaining how surviving SN520 PD-GSCs might have escaped apoptosis – by transitioning into the MES subtype (Supplementary Figure S4). Finally, a majority of Leiden clusters did upregulate genes associated with autophagy (Supplementary Table S2), which aligns with previously reported mechanisms of pitavastatin in glioma cells [5], and suggests that autophagy played a role in the response of SN520.

***TF-TF network modeling and ODE simulation motivation***

As characterization of drug response at single-cell resolution strongly supported the notion that the PD-GSCs underwent drug-induced transitions, we sought to understand how transcriptional regulatory mechanisms could govern the phenotypic heterogeneity observed within and across the two PD-GSC populations. Thus, we investigated the dynamical behavior of the underlying transcriptional regulatory networks from which multiple steady states, i.e., phenotypic states, can emerge. The TFs comprising each core network were all associated with response-relevant processes (Supplementary Table S1) [6–59]). Using the core TF-TF networks (Fig. 4E, F), we applied a previously developed algorithm known as random circuit perturbation (RACIPE, [60,61]), originally designed to model EMT circuits in cell development and other cancers [62–64]. Briefly, RACIPE generates an ensemble of ordinary differential equation (ODE) models based on associated chemical rate equations with distinct, random kinetic parameter sets. Because distinct, random kinetic parameters sets are used, this avoids the main issue of parameter identification for kinetic-based ODE models. From the ensemble of models, we would be able to analyze the resulting distribution of steady states and identify robust phenotypes supported by the core TF network. Previous applications have demonstrated that this ensemble modeling/simulation approach was able to recapitulate established cell states in the context of EMT-associated metastasis, B-cell lymphopoiesis, and small cell lung cancer [60,65–67].

***TF-TF network validation for SN520 and SN503***

To test the predictive capabilities of the TF-TF network topologies, we evaluated how similar or dissimilar the simulated states were to experimental data when each network was initialized using untreated (D0) data for each PD-GSC, respectively. Hierarchical clustering of the simulated steady states for both SN520 and SN503 resulted in four main clusters, i.e., "robust" steady states (Fig 6C, E – dendrogram of simulated states). Coincidentally, four main clusters were identified from the untreated (D0) and pitavastatin-treated SN520 and SN503 PD-GSCs as well. We then determined pairwise cosine similarity values derived from pairwise comparisons of *i)* PD-GSCs to one another within each hierarchical cluster and *ii)* simulated states to PD-GSCs within hierarchical clusters. Of the latter comparisons, when clusters of simulated states were similar to experimental clusters, the distributions of cosine similarity values were significantly higher than distributions based on comparisons in which TF gene expression was randomly permuted (Supplementary Figure S10).

To assess the statistical significance of the network topologies, we also performed simulations in which nodes and edges were randomly assigned such that the number of nodes and connections remained the same. Using the same untreated (D0) TF expression values as initial conditions, we performed RACIPE simulations using 1000 permutated networks, where each permuted network was used to run 1000 randomly selected parameter sets with a randomly selected untreated (D0) TF expression profile. The resulting 1e6 simulated states for each PD-GSC were then compared to the TF expression profiles of untreated and pitavastatin-treated cells to create a null distribution of cosine similarity values. Using this null distribution as a basis of comparison, we found that cosine similarity values derived from the original TF-TF network topologies were statistically significant (SN520 p-value $< 1e-16$, p-value SN503 $< 1e-16$, Supplementary Figure S10).

11. Pölönen P, Jawahar Deen A, Leinonen HM, et al. Nrf2 and SQSTM1/p62 jointly

contribute to mesenchymal transition and invasion in glioblastoma. *Oncogene*. 2019;38(50):7473-7490. doi:10.1038/s41388-019-0956-6

49. Wang B, Ma Q, Wang X, Guo K, Liu Z, Li G. TGIF1 overexpression promotes glioma progression and worsens patient prognosis. *Cancer Med*. 2022.

50. Weng C-C, Hsieh M-J, Wu C-C, et al. Loss of the transcriptional repressor TGIF1 results in enhanced Kras-driven development of pancreatic cancer. *Mol Cancer*. 2019;18(1):96. doi:10.1186/s12943-019-1023-1

51. Lorenz NI, Sittig ACM, Urban H, et al. Activating transcription factor 4 mediates adaptation of human glioblastoma cells to hypoxia and temozolomide. *Sci Rep*. 2021;11(1):14161. doi:10.1038/s41598-021-93663-1

52. Tseng Y-H, Huang Y-H, Lin T-K, et al. Thyroid hormone suppresses expression of stathmin and associated tumor growth in hepatocellular carcinoma. *Sci Rep*. 2016;6(1):38756. doi:10.1038/srep38756

53. Ruan X, Zhang R, Li R, et al. The Research Progress in Physiological and Pathological Functions of TRAF4. *Front Oncol*. 2022;12(February):1-13.

doi:10.3389/fonc.2022.842072

### Supplementary Figure Legends

**Figure S1. Pitavastatin-induced kill kinetics in SN520 and SN503**. Plots of mean cell viability during pitavastatin treatment for SN520 (top) and SN503 (bottom). Each series of relative viability values corresponds to a different pitavastatin concentration. Relative viability was calculated with respect to the untreated (vehicle-normalized, pitavastatin = 0.0μM) condition. Plotted values are mean viability values (N = 3) and error bars represent $\pm$ 2x standard deviation.

**Figure S2. Flow cytometry analysis of apoptosis and cell death.** Dot plots of cells assessing cell death (SYTOX) and apoptotic markers (annexin V) for **(A)** SN520 and **(B)** SN503. Gating was based on an unstained control and heat-inactivated/live cell (50:50) mixed control sample for each PD-GSC. Heat inactivation consisted of incubating cells in 60°C water bath for 15 minutes, with a small sample being inspected post incubation under microscope to ensure that inactivated cells were not completely lysed. Due to sample-limitations in D2 pitavastatin treatment, flow cytometry assessment of cell-death and apoptosis was not performed (N/A plot).

**Figure S3. Defining GBM molecular subtypes via gene expression.** Heatmaps of subset of 20 genes used to define GBM molecular subtypes for **(A)** SN520 and **(B)** SN503 cells. Violin/boxplots of GSVA enrichment scores (ES) for CL, PN, and MES molecular subtypes determined for cells. Clusters of violin/boxplots correspond to molecular subtype scores for cells categorized to each subtype for **(C)** SN520 and **(D)** SN503. Those cells having a negative ES for all three subtypes remained undefined (TBD - grey).

**Figure S4. Cell cycle phase breakdown of SN520 and SN503. (A)** UMAP plot of SN520, similar to Figure 2A, annotated for cell cycle phase for each cell. **(B)** Proportions of cells in each cell cycle phase within each treatment condition for SN520. **(C)** UMAP plot of SN503 annotated for cell cycle phase for each cell. **(D)** Proportions of cells in each cell cycle phase within each treatment condition for SN503.

**Figure S5. DNA quantification throughout treatments.** Density plots of fluorescent signals generated from cells stained with propidium iodide (PI) throughout pitavastatin- (top) and vehicle-treatment (bottom) for **(A)** SN520 and **(B)** SN503. Portions of the density plots representative of specific cell cycle phases have been labeled, along with percentages of cells within each phase.

**Figure S6. Theoretical calculations corroborate PMT rather than selection. (A)** Summary of cell counts used to determine doubling times ($t_d$) for SN520 and SN503. **(B)** Calculations supporting three scenarios that potentially explain the increase in the proportion of MES cells within SN520. Scenario *i*) assumes a selection of pre-existing MES PD-GSCs. Scenario *ii*) assumes exponential growth of MES cells only with a $t_d$ based on (A). Finally, scenario *iii)* assumes exponential growth of MES cells, but determines a $t_d$ that would enable MES growth to match final SN520 cell counts on the fourth day of pitavastatin treatment. The corresponding $t_d$ required to achieve a final MES cell count is listed.

**Figure S7. DEG and Cell cycle phase proportions within cell clusters.** (**A)** Heatmap of the top upregulated DEGs, based on FDR values, across the Louvain cell clusters (cl) identified in vehicle-control- and pitavastatin-treated cells for **(A)** SN520 and **(B)** SN503. Labeled genes are representative members of various enriched hallmark gene sets (CH = cholesterol homeostasis, OP = oxidative phosphorylation, MTOR = MTORC1 signaling, EMT = epithelial-to-mesenchymal transition). **(C-D)** Proportions of cells in each cell cycle phase within each Louvain cluster for SN520 and SN503, respectively.

**Figure S8. SN520 and SN503 regulon activities.** Heatmaps of eigengene values, i.e., regulon activities for **(A)** SN520 and **(B)** SN503. Top row of heatmaps show regulon activities in cells rearranged according to corresponding pseudotime. Bottom row of heatmaps include cells rearranged with respect to experimental treatment. Top color bars represent pseudotime, treatment, and MINER3-inferred transcriptional state.

**Figure S9. Transcriptional program activity dynamics.** Activity profiles of remaining transcriptional programs not included in main Figure 5D for **(A)** SN520 and **(B)** SN503. Programs were clustered together based on their LOESS regressed activity profiles with respect to pseudotime. Dashed grey lines represent the average shape of regression profiles for each program cluster. Representative hallmark gene sets enriched within programs are included within each plot.

**Figure S10. Significance and validation of TF-TF network topologies. (A)** Boxplots of pairwise cosine similarity values based on specific pairwise comparisons of experimental data and simulated states generated from SYGNAL-520. Horizontal lines with adjacent asterisks connecting two distributions indicate that the first (leftmost) distribution is statistically significantly higher (FDR < 0.01). Distribution of cosine similarity values from pairwise comparisons of experimental states ($ES_{520}$) and permutated states are shown in grey. Permutated states were derived from *i)* permuted expression data, where both cell and gene labels were randomly permuted, and *ii)* randomized network topology (METHODS). **(B)** Boxplots for SYGNAL-503. Note that multiple simulated states ($SS_{503}$-2 and $SS_{503}$-4) showed similarities to $ES_{503}$-3. **(C)** Heatmap of mean relative expression (z-score) of TFs across cells within each experimental state (ES) and simulated state (SS) for SN520, states (columns) are hierarchically clustered. Color bars on top indicate states and data type (grey – simulation or black – experimental data). PCA plot of simulated states from SYGNAL-520 below the heatmap is included for reference. **(D)** Corresponding heatmap of mean relative expression (z-score) of TFs and PCA plot for SN503. **(E)** Heatmap (left) of mean relative expression for TFs in experimental states and subset of simulations from SYGNAL-503 in which the input expression value for ARID5A, MEOX2, and MAFF are high (>1 normalized expression). States (columns) are hierarchically clustered. Color bar above indicates cell states being compared. Adjacent heatmap (right) shows pairwise cosine similarity values of mean relative expression profiles of experimental and simulated states that have high levels of ARID5A, MEOX2, and MAFF. Color bars indicate cell states. **(F)** Dot plot of TFs rank-ordered based on their importance in classifying experimental states for SN503 using random forest analysis.

**Figure S11. Convergence of RACIPE simulations of TF-TF networks. (A)** Kullback-Leibler divergence distances (black) of simulated states generated by SYGNAL-520 with respect to one another. Simulated states were generated using the respective TF-TF network using a different number of model simulation parameters (1e3, 2e3, 4e3, 6e3, 8e3, and 1e4 randomly selected model parameters) across 100 randomly selected initial conditions. Number of unique states (blue) is based on the number of steady states identified having a specified Euclidean distance greater than its nearest neighbors (METHODS). **(B)** Kullback-Leibler divergence distances normalized with respect to number of unique states identified per set of simulations performed using SYGNAL-520. **(C-D)** Kullback-Leibler divergence distances and normalized distances, respectively, determined from simulations using SYGNAL-503.

**Figure S12. Random forest model predicts cell state with high-level of accuracy. (A)** PCA plot of a randomly selected subset of 2,000 simulated states, i.e., TF expression profile, from the 1 million simulations performed using SYGNAL-520 (Fig 6C), with each dot representing a simulation output. Fill colors for each dot represent the state of the cell as defined by hierarchical clustering of the cells. Border colors represent the state of the predicted simulated state using the random forest model trained on a subset of the 1 million simulations. **(B)** Proportion of each actual state, defined by hierarchical clustering, within the predicted simulated states for SN520. **(C)** PCA plot of a randomly selected subset of 2,000 simulated states for SN503. **(D)** Proportion of actual states within each predicted state for SN503.

**Figure S13. *In silico* KD simulations in SYGNAL-520/503. (A)** Stacked bar plot depicting proportion of simulated states assigned to one of four simulated states ($SS_{520}$-i) identified from hierarchical clustering of RACIPE simulations using SYGNAL-520 (Figure 6C) in response to a 95% knock down (KD) in expression of particular TF. TFs are rank ordered according to the proportion of simulated states assigned to $SS_{520}$-1, which corresponds to a mesenchymal state. **(B)** Stacked bar plot depicting proportion of simulated states assigned to one of four simulated states ($SS_{503}$-i) identified from hierarchical clustering of RACIPE simulations using SYGNAL-503 (Figure 6E) in response to a 95% knock down (KD) in expression of particular TF. TFs are rank ordered according to the proportion of simulated states assigned to $SS_{503}$-1, which contained the largest proportion of mesenchymal cells.

**Figure S14. Pitavastatin pretreatment (48hrs) improves vinflunine efficacy in PD-GSCs. (A)** Dose-response curves for SN520 (top) and SN503 (bottom) pretreated with pitavastatin for 48 hrs and subsequently treated with vinflunine. Colors indicate specific pre-treatment conditions. Asterisks indicate statistically significant differences in relative viability across conditions. **(B)** Vinflunine $IC_{50}$ values measured across additional non-responder and responder PD-GSCs. Error bars represent 2x standard deviation. Color annotation identical to (A).

**Figure S15. SN520 exhibits a sequence of TF expression distinct from previously proposed mechanisms of PMT. (A)** Master regulators driving PMT in GBM. Figure modified from Fedele et al. 2019. **(B)** Sequence of TF expression of master regulators (Fedele et al. 2019) observed in SN520. Density plot and heatmap align with pseudotime (bottom color bar). Density plot shows proportion of cells belonging to each treatment condition arranged according to pseudotime. Heatmap shows relative expression (LOESS regression) of TF expression along pseudotime. TFs listed are master regulators overlapping in SN520 scRNA-seq data set. TFs are listed in sequential order according to

### Supplementary Figure S1

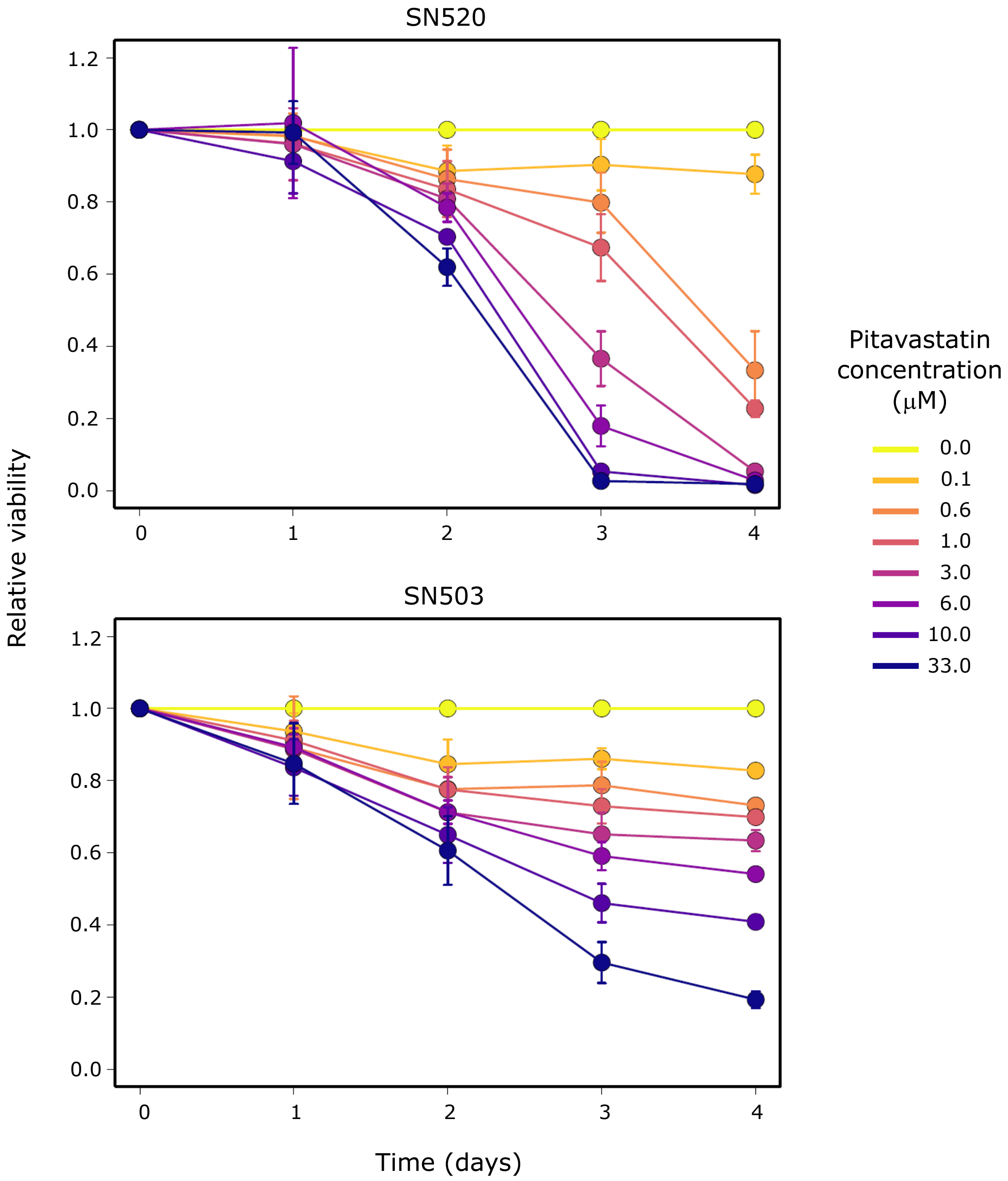

Supplementary Figure S2

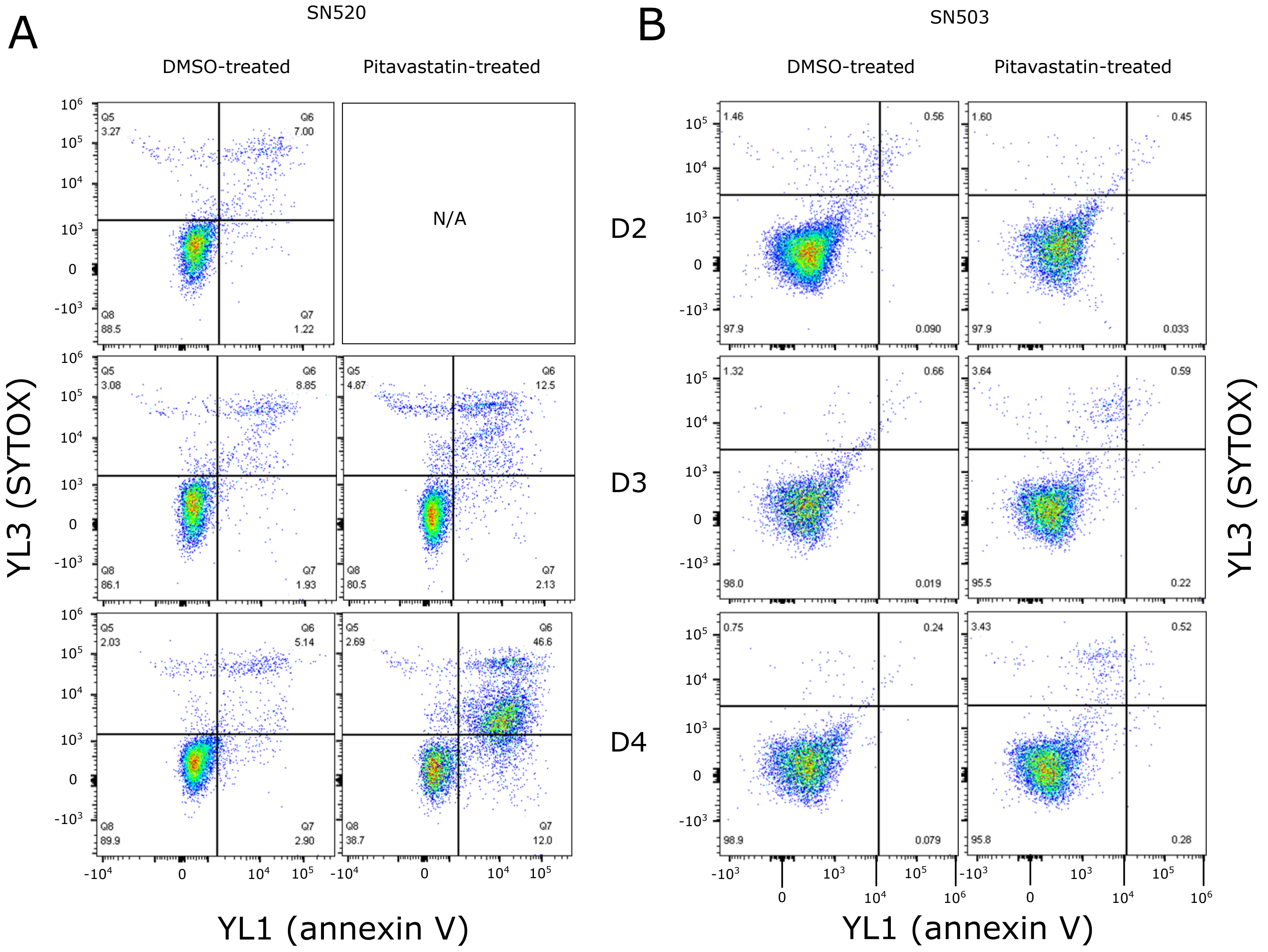

Supplementary Figure S3

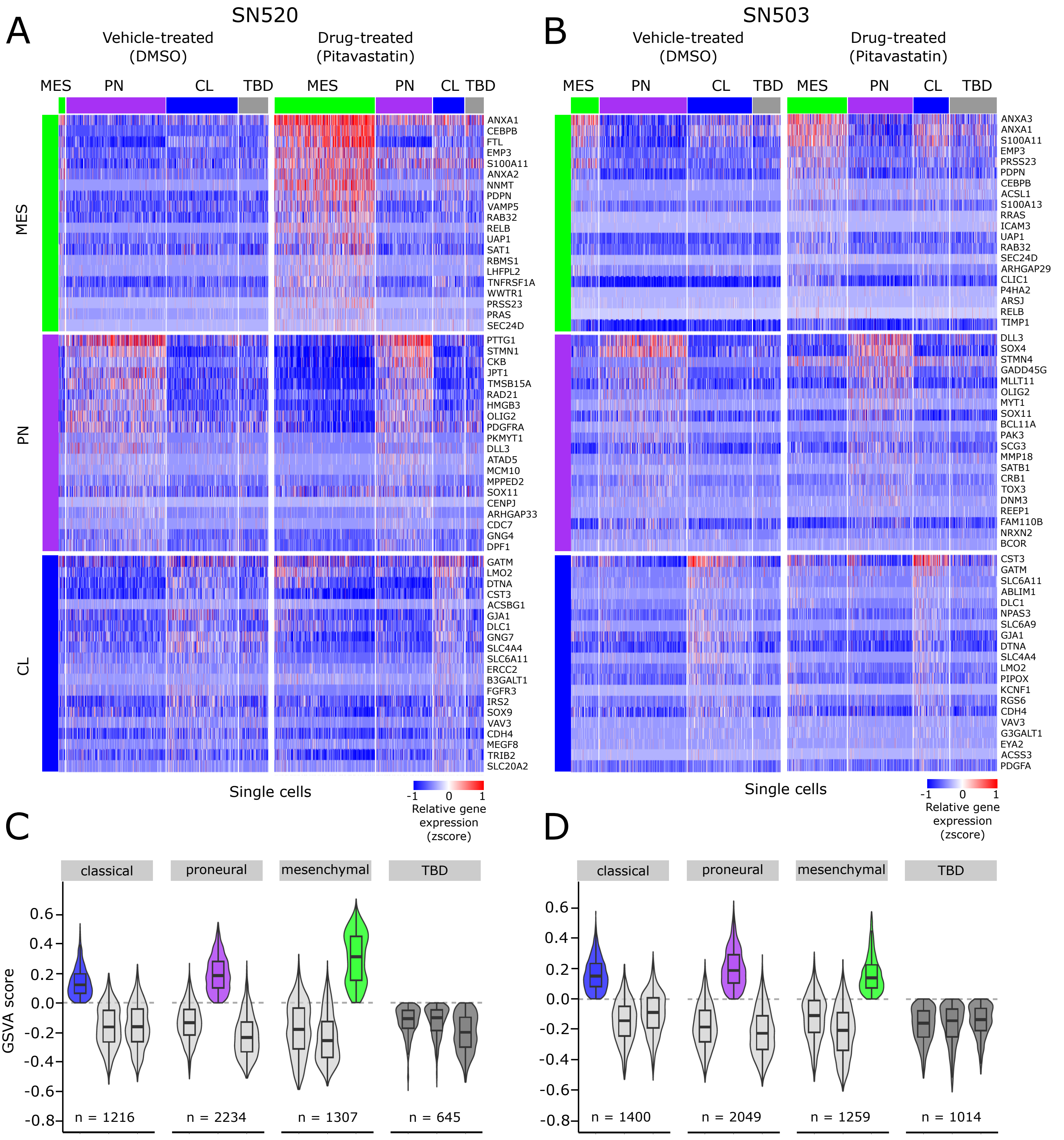

### Supplementary Figure S4 - Cell-cycle phase breakdown in treatments

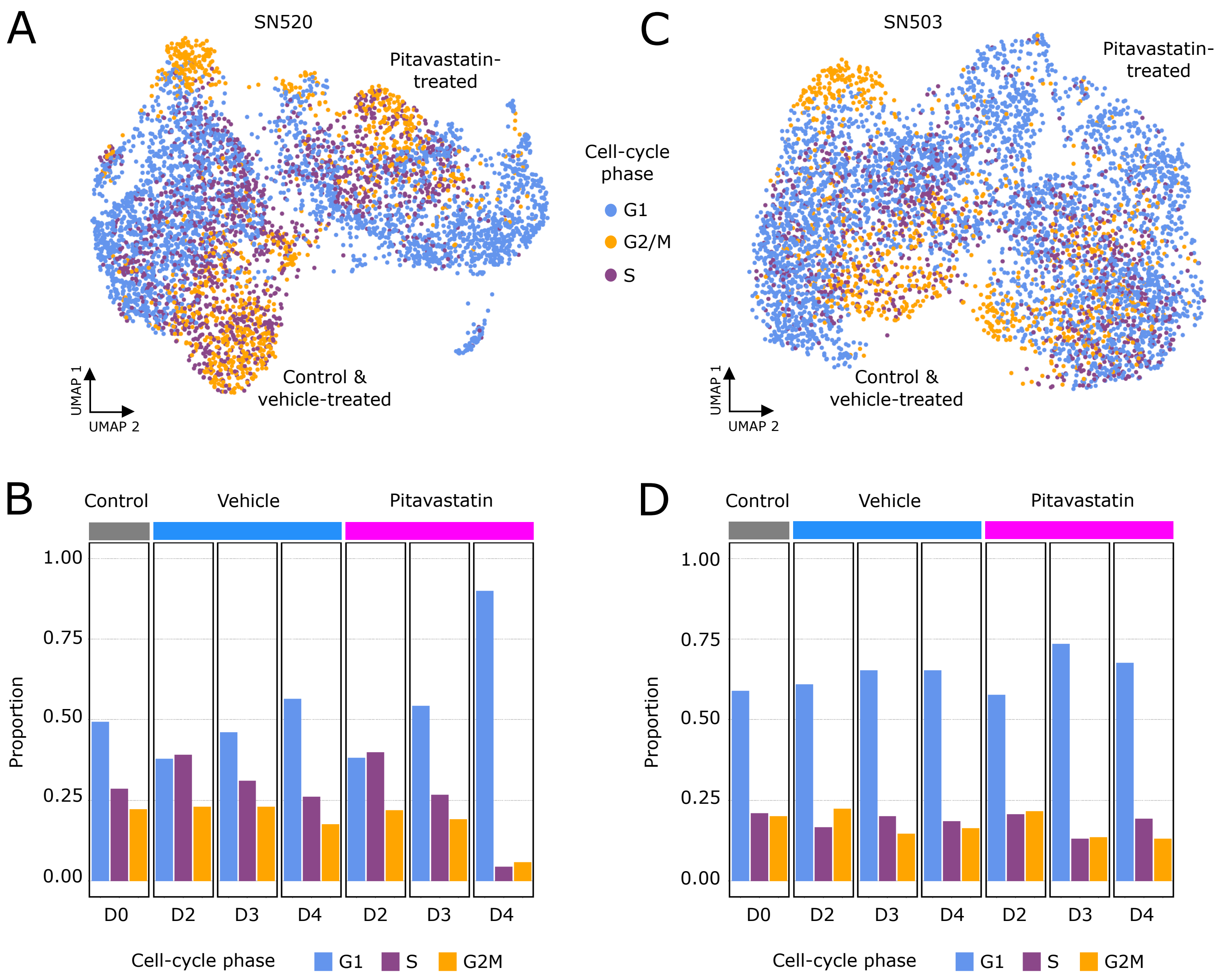

Supplementary Figure S5

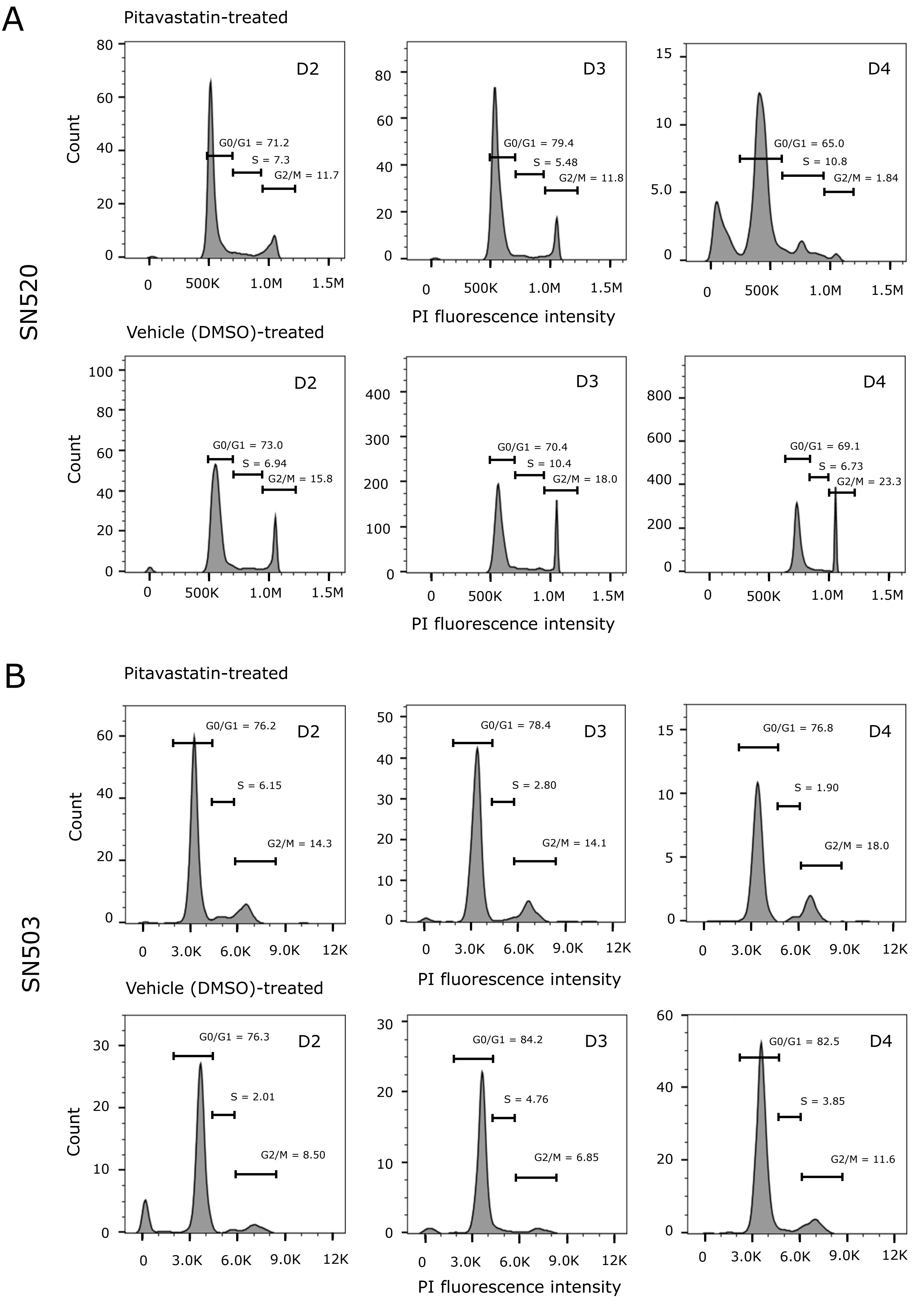

Supplementary Figure S6

A

| PD-GSC ID | Incoulation density (cells) | Passage/harvest density (cells) | Growth duration (days) | Substrate | Estimted $t_d$ (hrs) |
|---|---|---|---|---|---|
| SN520 | 4.36E+05 | 7.56E+06 | 14 | T75 flask | 81.62 |
| SN520 | 4.36E+05 | 7.48E+06 | 14 | T75 flask | 81.93 |
| SN520 | 4.36E+05 | 5.94E+06 | 14 | T75 flask | 89.17 |
| SN503 | 5.68E+05 | 5.58E+06 | 13 | T75 flask | 94.65 |
| SN503 | 5.68E+05 | 4.22E+06 | 13 | T75 flask | 107.79 |
| SN503 | 5.68E+05 | 4.40E+06 | 13 | T75 flask | 105.59 |

| | mean $t_d$ (hrs) |
|---|---|
| SN520 | 84.24 |
| SN503 | 102.68 |

B

**Scenario i**

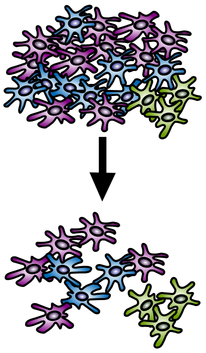

| Time (day) | Total cell count | MES fraction | MES cell count |
|---|---|---|---|
| 0 | 1.200E+06 | 0.021 | 2.466E+04 |

constant number of MES cells

| | | | |
|---|---|---|---|
| 4 | 5.375E+05 | 0.046 | 2.466E+04 |

**Scenario ii**

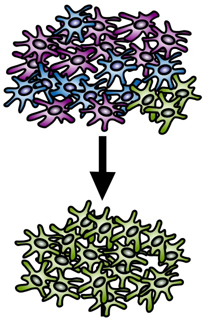

| Time (day) | Total cell count | MES fraction | MES cell count |
|---|---|---|---|
| 0 | 1.200E+06 | 0.021 | 2.466E+04 |

$x_f = x_i * 2^n$

$n = \Delta t / t_d$

$t_d = 84.24$ hrs (SN520)

| | | | |
|---|---|---|---|
| 4 | 5.407E+04 | 1.000 | 5.40E+04 |

**Scenario iii**

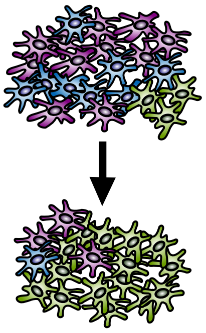

| Time (day) | Total cell count | MES fraction | MES cell count |
|---|---|---|---|
| 0 | 1.200E+06 | 0.021 | 2.466E+04 |
| 4 | 5.375E+05 | 0.940 | 5.054E+05 |

required $t_d = \Delta t / (\log 2(x_f/x_i))$

required $t_d = 21$ hrs

original $t_d = 21$ hrs

Proneural 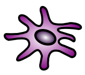 Classical 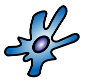 Mesenchymal 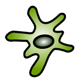

Supplementary Figure S7

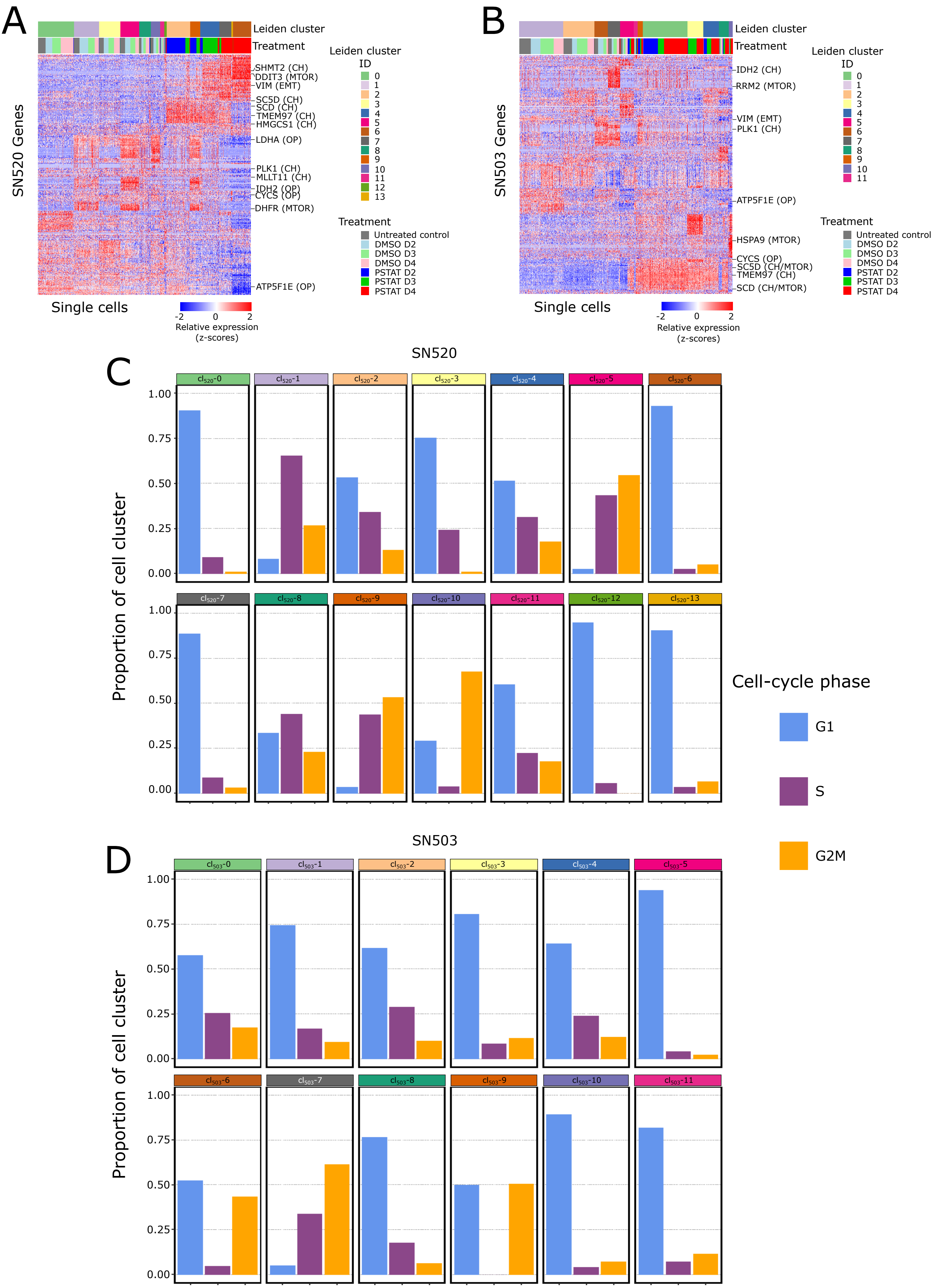

### Supplementary Figure S8

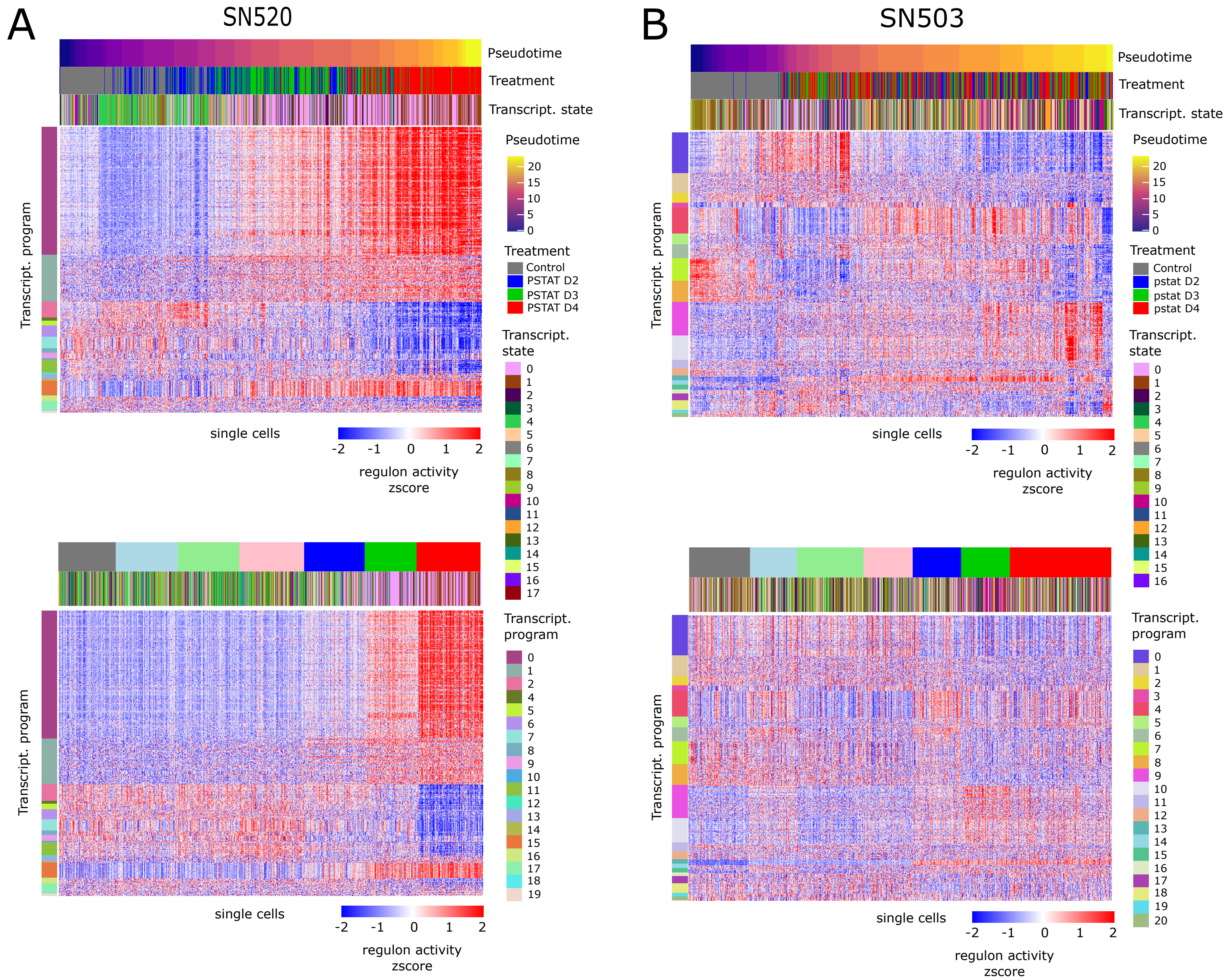

Supplementary Figure S9

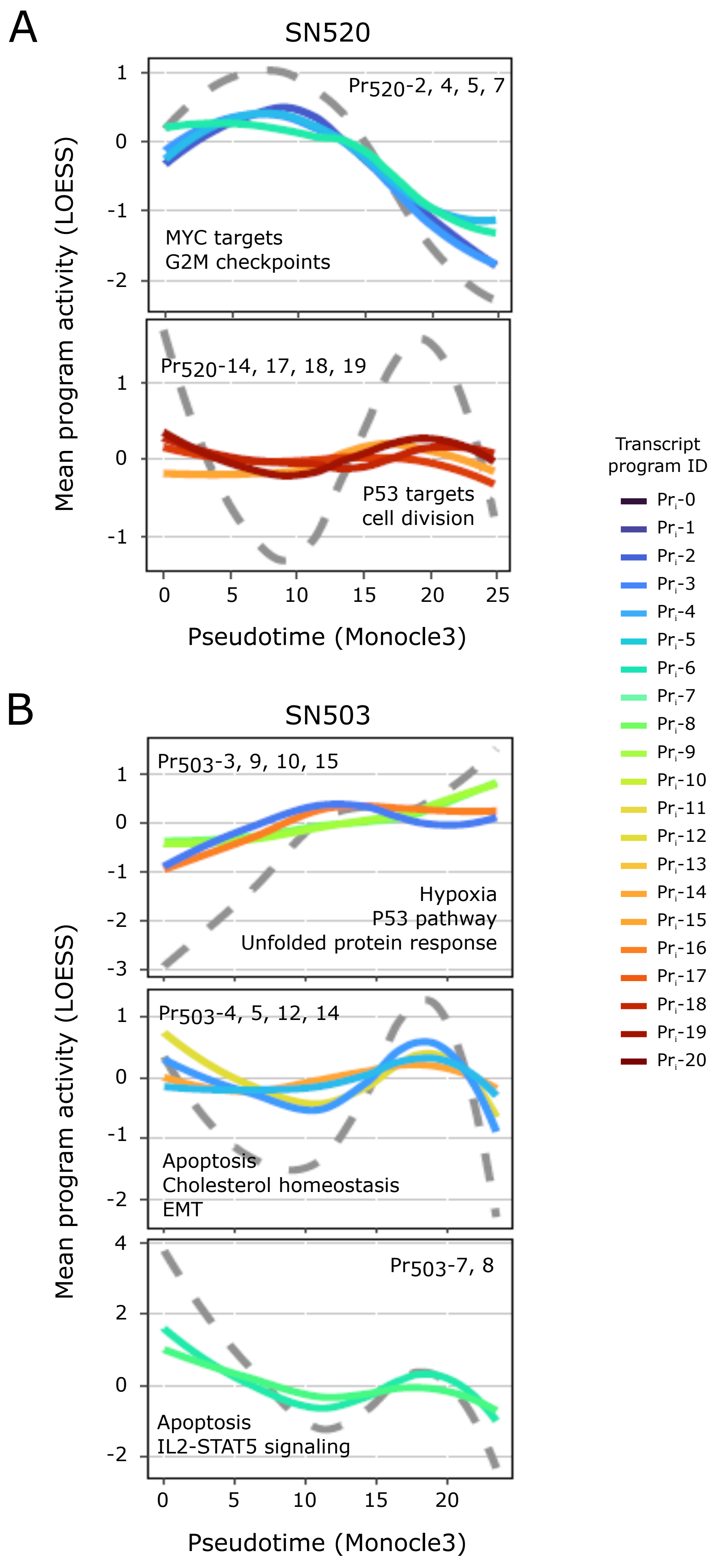

### Supplementary Figure S10

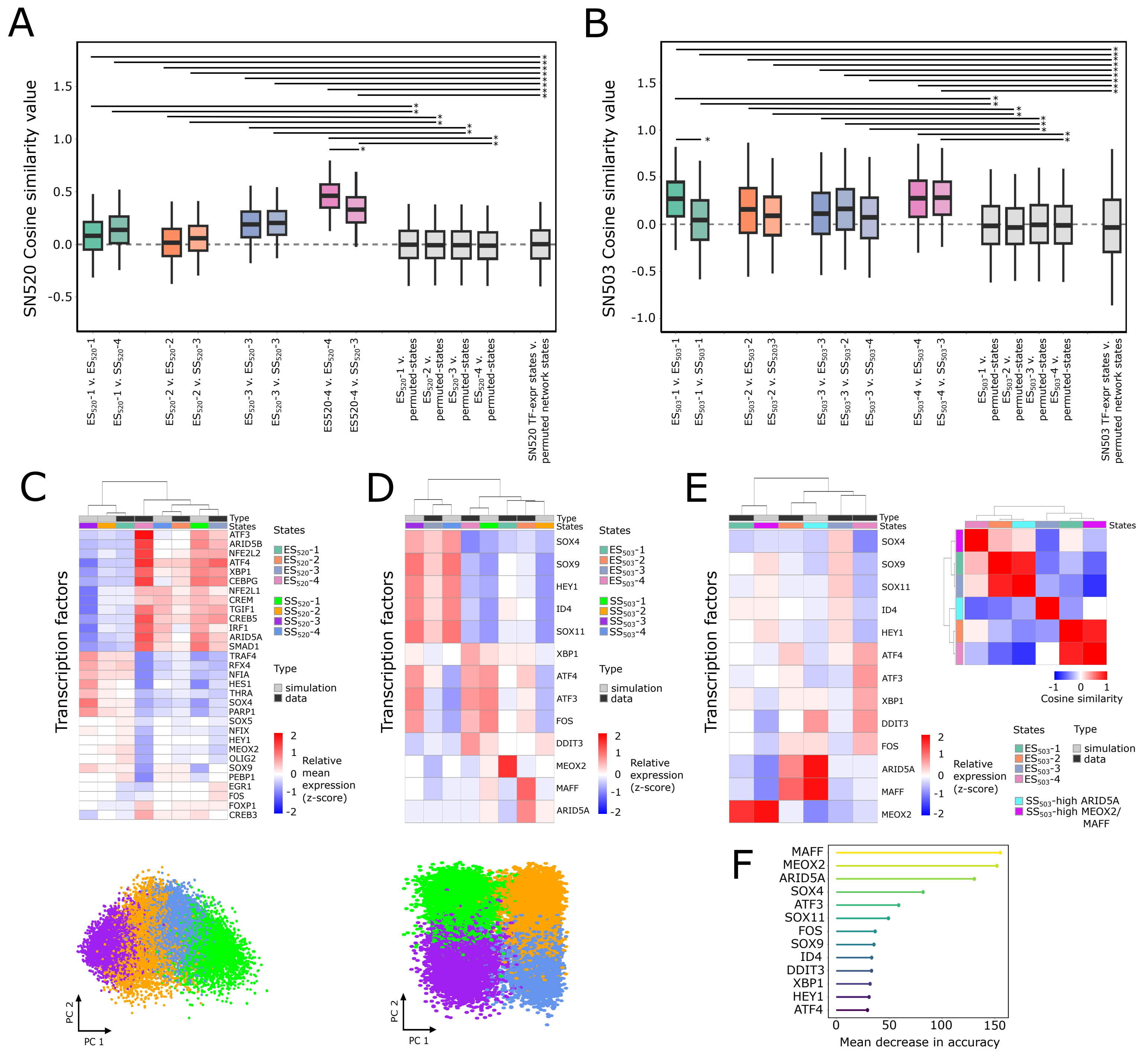

### Supplementary Figure S11

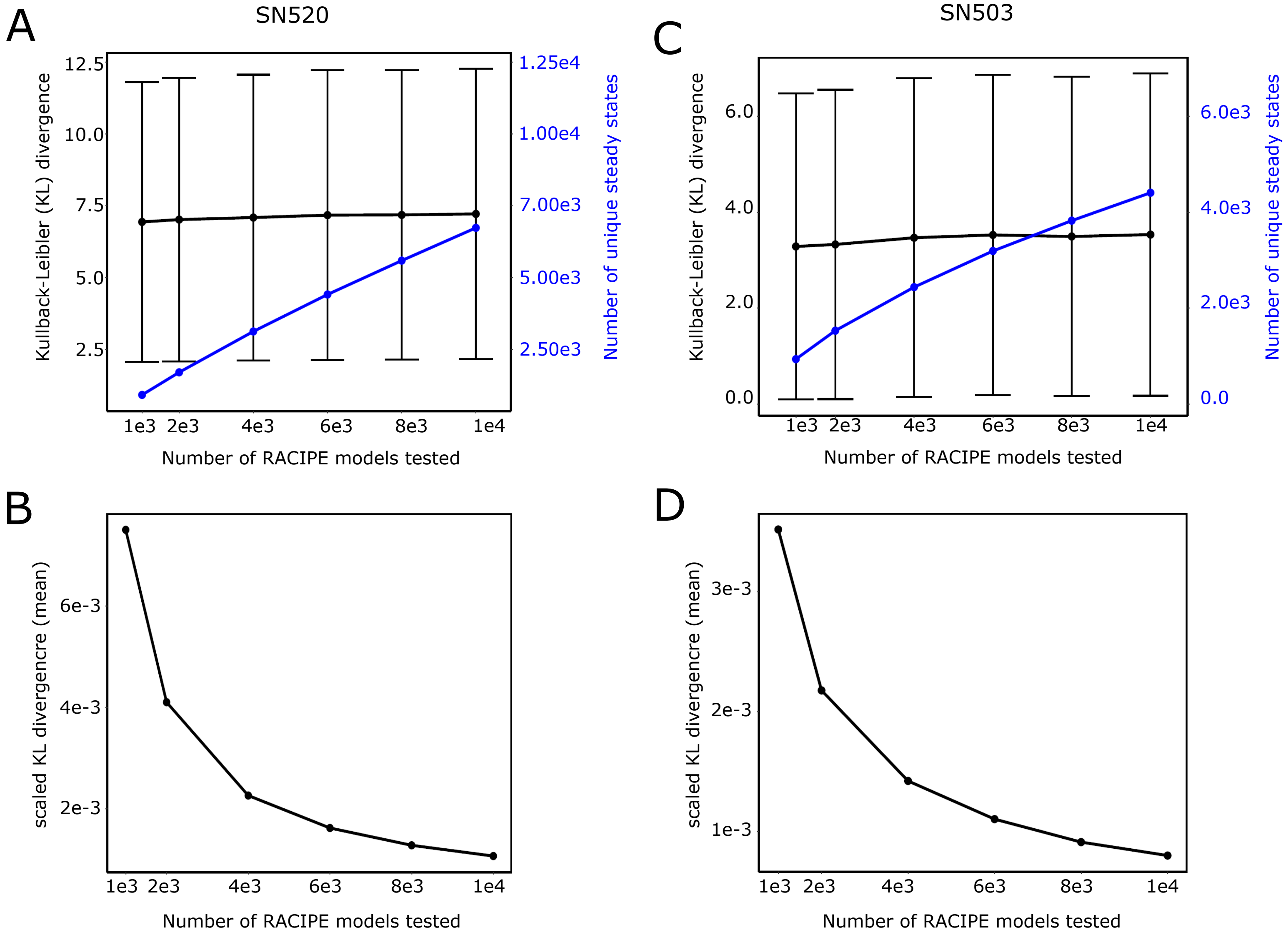

Supplementary Figure S12

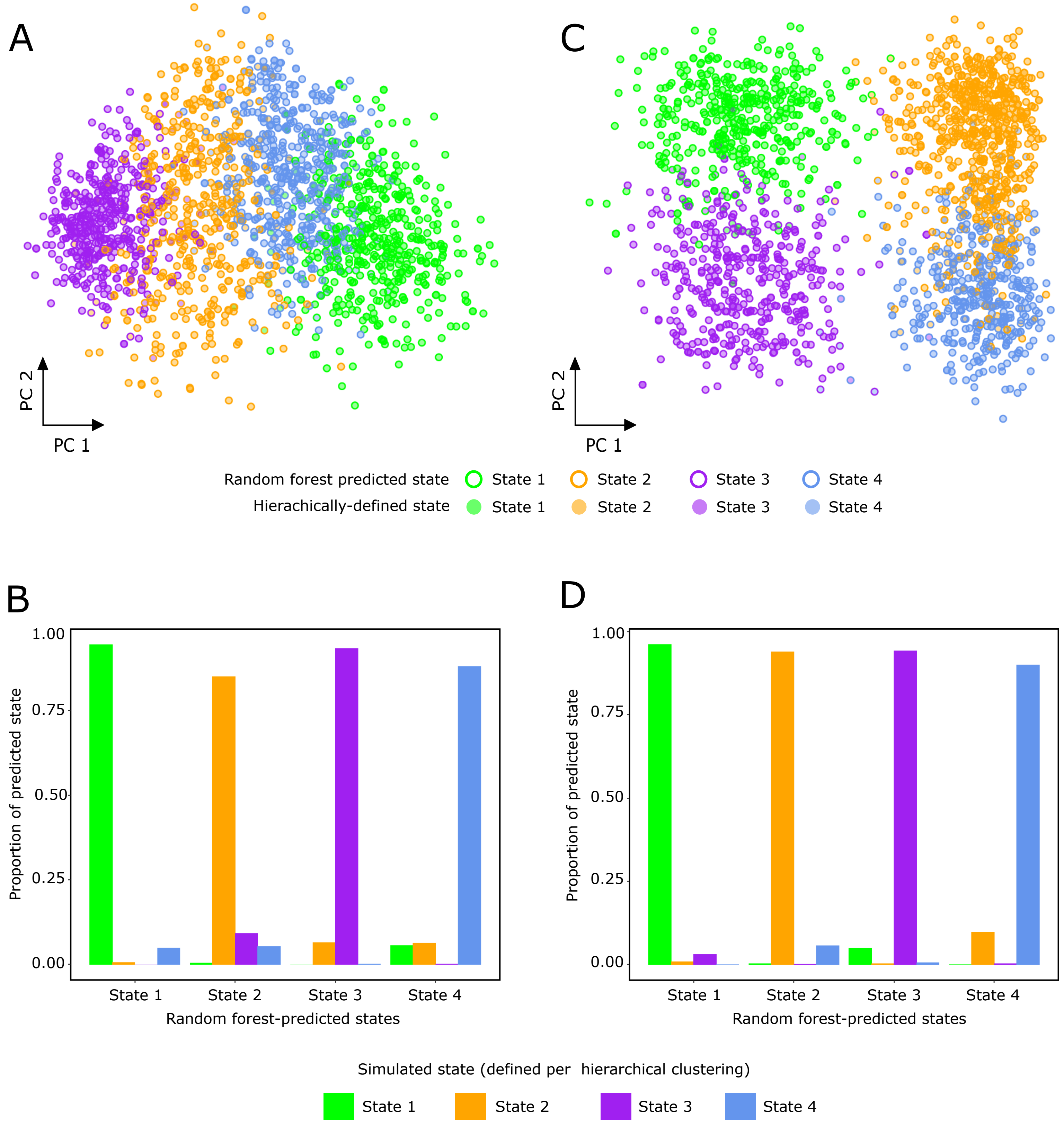

### Supplementary Figure S13

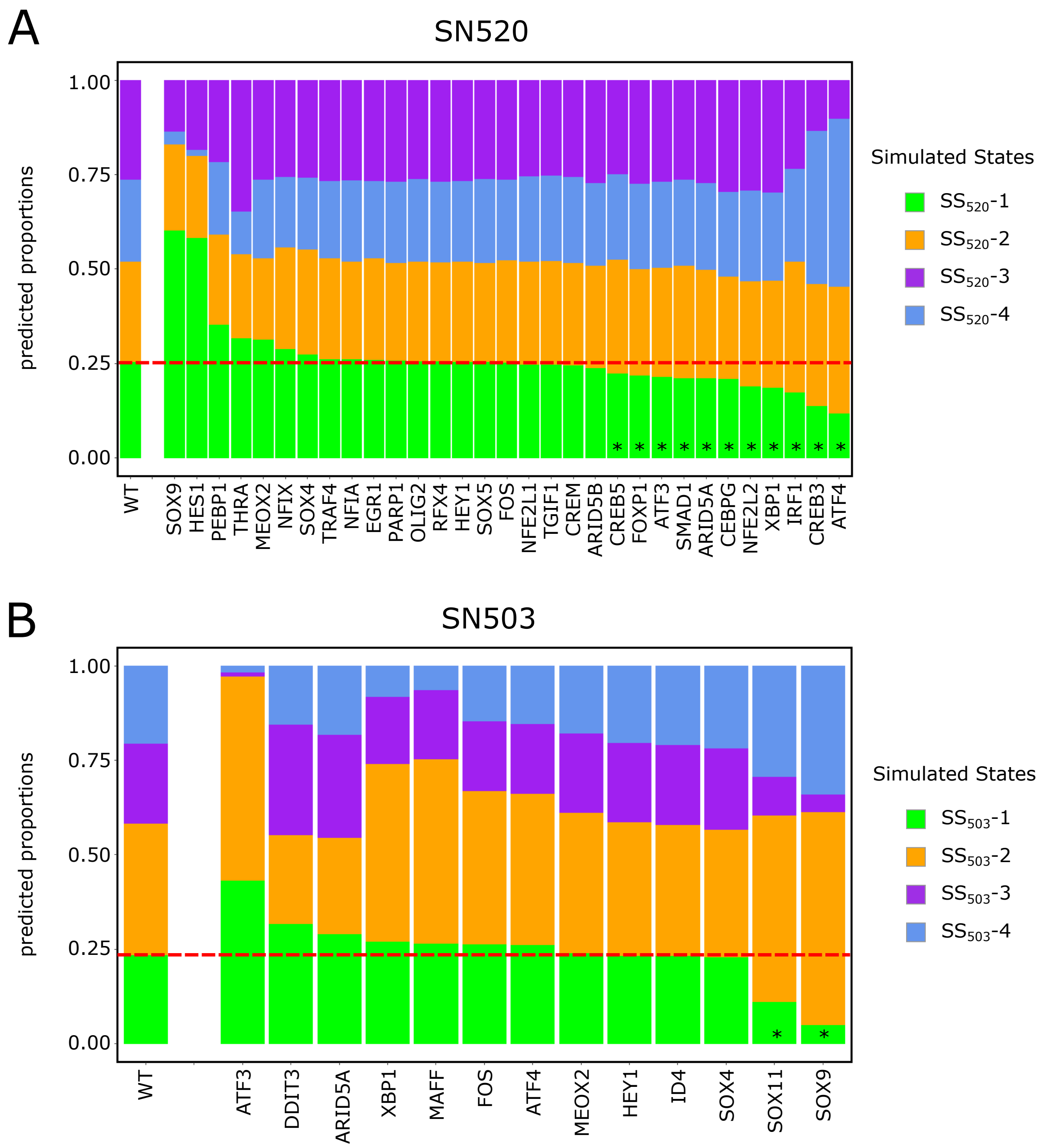

### Supplementary Figure S14

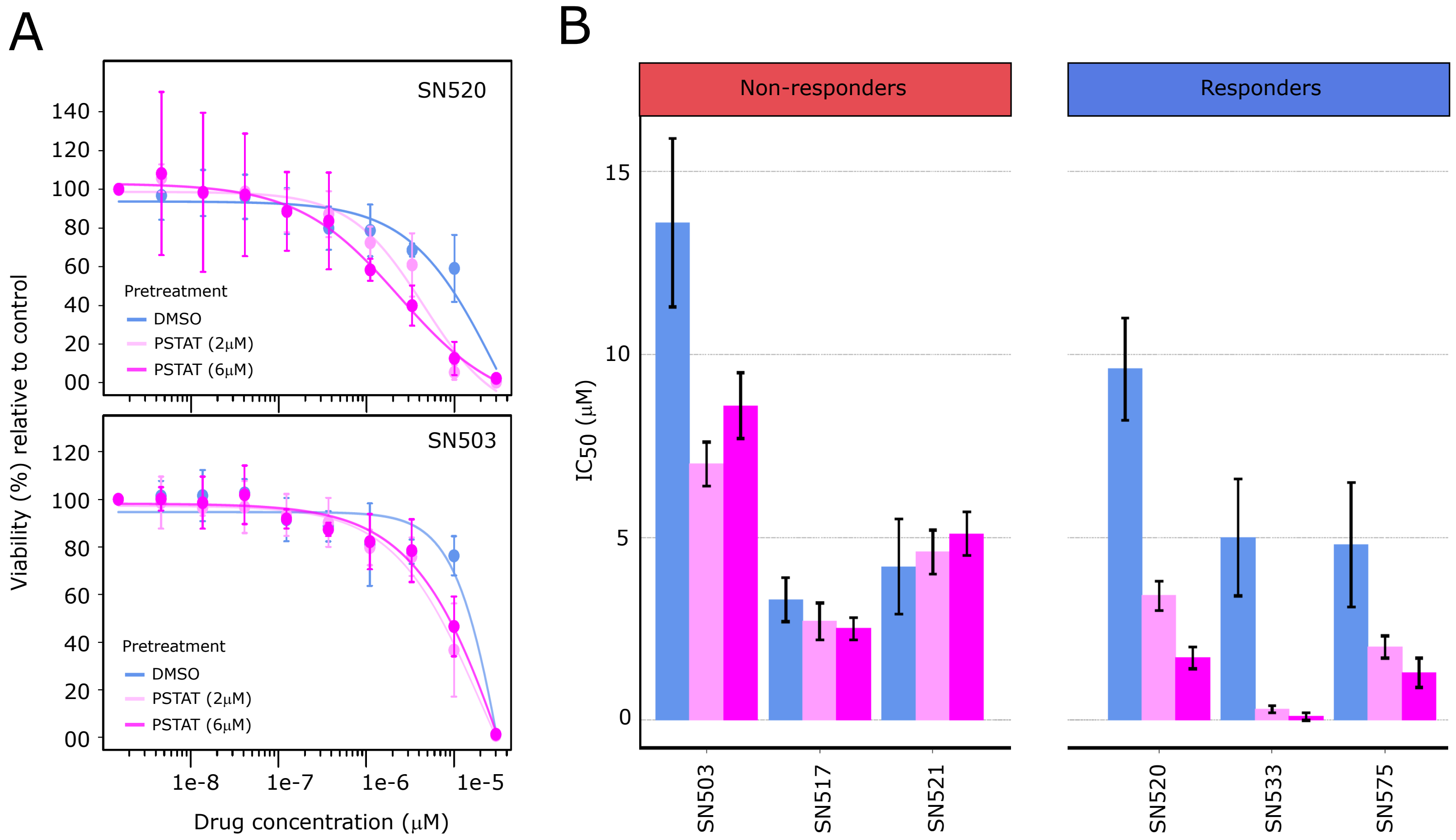

### Supplementary Figure S15

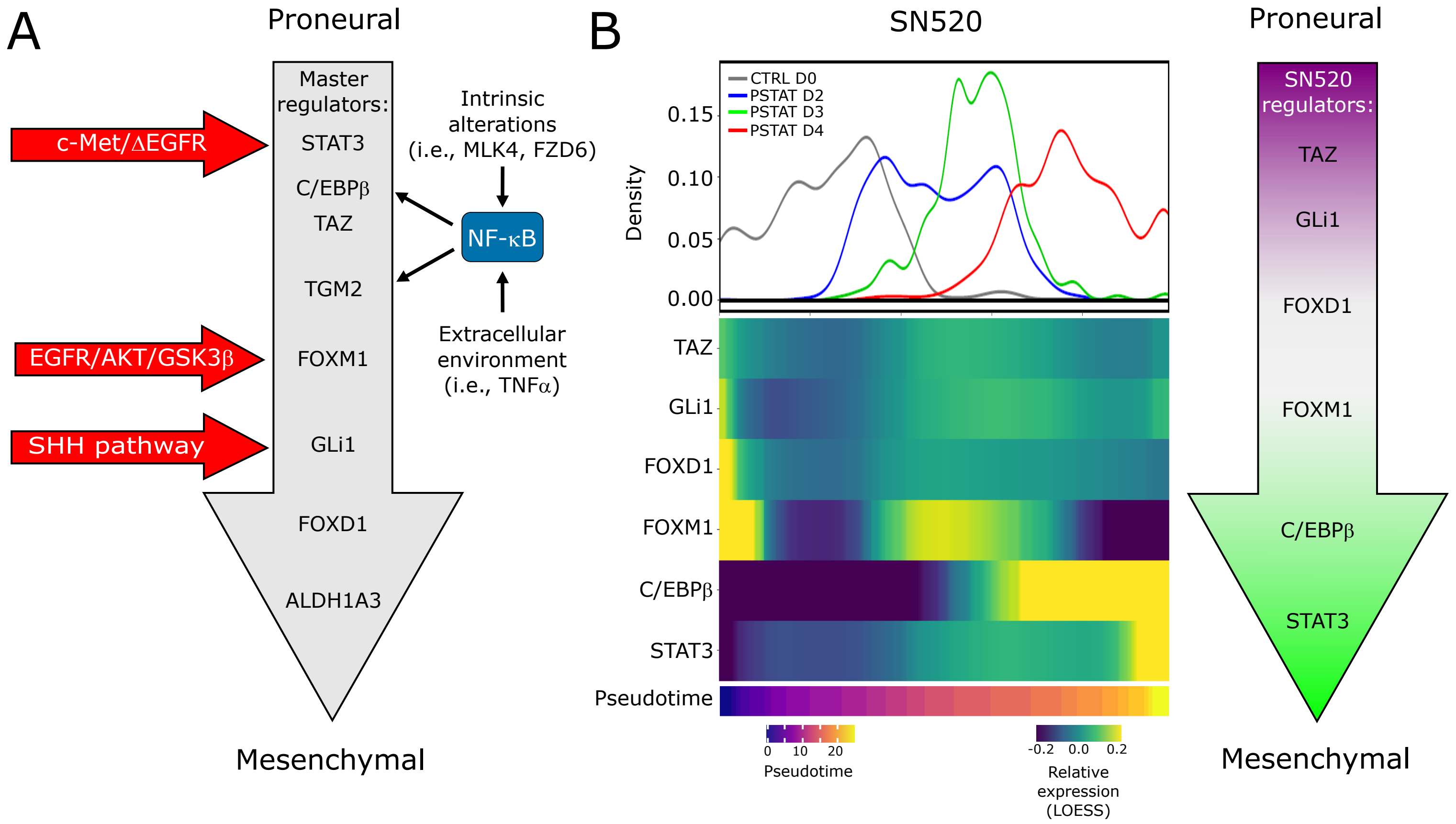
